# Non-functional and weak alleles of *FRIGIDA* and *FLOWERING LOCUS C* reduce lifetime water-use independent of leaf-level water-use-efficiency traits in *Arabidopsis thaliana*

**DOI:** 10.1101/443424

**Authors:** J.N. Ferguson, R.C. Meyer, K.D. Edwards, M. Humphry, O. Brendel, U. Bechtold

## Abstract

Natural selection driven by water availability has resulted in considerable variation for traits associated with drought tolerance and leaf level water-use efficiency (*WUE*). In Arabidopsis, little is known about the variation of whole-plant water use (PWU) and whole-plant *WUE* (TE). To investigate the genetic basis of PWU, we developed a novel proxy trait by combining flowering time and rosette water use to estimate lifetime PWU. We validated its usefulness for large scale screening of mapping populations in a subset of ecotypes. This parameter subsequently facilitated the screening of water-use but also drought tolerance traits in a recombinant inbred line population derived from two Arabidopsis accessions with distinct water use strategies, namely C24 (low PWU) and Col-0 (high PWU). Subsequent quantitative trait loci (QTL) mapping and validation through near-isogenic lines identified two causal QTLs, which showed that a combination of weak and non-functional alleles of the *FRIGIDA (FRI)* and *FLOWERING LOCUS C (FLC)* genes substantially reduced plant water-use without penalising reproductive performance. Drought tolerance traits, stomatal conductance, intrinsic water use efficiency (δ^13^C) and rosette water-use were independent of allelic variation at *FRI* and *FLC*, suggesting that flowering is critical in determining life-time plant water use, but not leaf-level traits.

## Introduction

Water availability is essential for the optimal allocation of resources to achieve maximal growth and reproductive fitness (Anderson 2016). Consequently, a water deficit may force survival trade-off costs resulting in reduced reproductive fitness (Von Euler, Ågren & Ehrlén 2014; Sletvold & Ågren 2015). In natural populations, adaptations to water deficits encompass a number of different ecological strategies that include drought escape and avoidance leading to drought resistance. While drought escape is characterised by rapid growth and early flowering to reproduce before the onset of terminal drought, avoidance limits growth during periods of dehydration through lowering stomatal conductance and transpiration (Ludlow 1989; Kooyers 2015). Drought resistance traits, characterised by the ability to survive a water deficit, have traditionally been used to assess plant performance under reduced water availability. However, the usefulness of drought resistance as a trait to optimize plant productivity has been questioned, as the improvement of various drought resistance related traits has been demonstrated to reduce productivity under some circumstances, regardless of the ability of plants to survive the period of drought stress (Blum 2005, 2009; Passioura 2007). It is widely accepted that drought resistance facilitates plant survival, but it does not contribute towards the maintenance of yield following drought stress or in water replete conditions (Blum 2005, 2009; Passioura 2007). The identification of plant varieties that are able to produce stabilized or improved yields with reduced water inputs is therefore an important goal for plant breeders, physiologists and molecular biologists alike (Parry, Flexas & Medrano 2005; Morison, Baker, Mullineaux & Davies 2008).

Water use efficiency at the leaf level is the net amount of CO_2_ fixed per unit of transpired water, hereafter referred to as instantaneous water-use-efficiency (*WUEi*, *A/E*) (Condon *et al*. 2004; Table 1). It relates equally to water loss by transpiration and net carbon gain achieved via gas exchange (Long, Marshall-Colon & Zhu 2015). Alternatively, carbon isotope composition (δ^13^C; Table 1), as an estimator of intrinsic water use efficiency, that is the ratio of net CO_2_ assimilation to stomatal conductance for water vapour (*A*/*g_s_*; Farquhar & Von Caemmerer 1982; Farquhar *et al*. 1989), are regularly used to describe integrated leaf level intrinsic water use efficiency and have been targeted in several studies as a primary trait to achieve “more crop per drop” as well as enhancing drought resistance (Morison *et al*. 2008; Blum 2009).

**Table 1:** Glossary of water use efficiency and water use parameters.

| Parameter | Abbreviation | Calculations |
| --- | --- | --- |
| Carbon isotope composition | $\delta^{13}\text{C}$ | $\frac{^{13}\text{C}}{^{12}\text{C}}$ |
| instantaneous leaf level water use efficiency | $WUE_i$ | $\frac{A}{E}$ |
| absolute vegetative (rosette) water use | VWU | slope 1 of linear regression<br>$slope = \frac{rSWC}{day} - intercept$ |
| calculated plant water use | cPWU | $VWU * daysofflowering$ |
| measured plant water use | mPWU | $\sum dailyaddedwater$ |
| mean daily water use | - | average of daily added water over the life time of the plant |
| water productivity calculated or measured | cWP / mWP | $\frac{seedbiomass}{cPWU \vee mPWU}$ |
| transpiration efficiency calculated or measured | cTE/ mTE | $\frac{abovegroundbiomass}{cPWU \vee mPWU}$ |
| dehydration plasticity (VWU plasticity) | DP | segmented regression<br>$\frac{(slope1 - slope2)}{slope1}$ |

The value of leaf level water use efficiency estimates for improving crop yield has previously been questioned. For example, it has been shown that despite the association between δ^13^C and water use efficiency in many species (Farquhar *et al*. 1989), its relation to yield across multiple environments and genotypes is much often variable (Condon *et al*. 2004). This suggests that both additional intrinsic plant factors, as well as environmental conditions, impact the relationship between intrinsic water use efficiency and agronomic water use efficiency, i.e. the amount of yield produced per unit of water transpired. Therefore, leaf level intrinsic water use efficiency estimates may not be a useful proxy to select for yield under water limited conditions. This lack of consistent upscaling from leaf-to whole-plant water use efficiencies may be a product of the heterogeneity of net CO_2_ assimilation rates within and across individual photosynthetic organs or it may also be due in part to the lack of integration of night-time transpiration and plant respiration rates in leaf-level WUE measurements (reviewed in Cernusak, Winter & Turner 2009; Cernusak *et al*. 2013). Furthermore, this inconsistency may be related to changes in environmental conditions leading to variations in other processes that affect CO_2_ supply and demand (Seibt, Rajabi, Griffiths & Berry 2008; Medrano *et al*. 2015). In addition, discrepancies may occur due to genotypic variation in carbon isotope signatures of crop plants being often driven by variation in stomatal conductance (Blum 2005; Monclus *et al*. 2006; Monneveux, Sánchez, Beck & Edmeades 2006; Marguerit *et al*. 2014), thereby limiting carbon assimilation and productivity. It should be noted, however, that in some species variation in δ^13^C has also been attributed to variation in carbon fixation as well as stomatal conductance (Masle, Gilmore & Farquhar 2005; Donovan, Dudley, Rosenthal & Ludwig 2007; Brendel *et al*. 2008).

Investigating the natural variation in whole-plant water-use efficiency and the mechanisms of drought resistance in natural populations is challenging, due to difficulties in re-creating realistic drought conditions in an experimental setting. For example, in short-dehydration experiments (Bechtold *et al*. 2010, 2016; Ferguson *et al* 2018), water loss is greater in larger plants creating substantial heterogeneity in the timing of water deficits (Kooyers 2015). While plant size greatly contributes to water use in Arabidopsis, drought response traits are independent of the transpiring leaf surface (Ferguson *et al*. 2018). This suggests that above ground biomass impacts water use and consequently whole-plant water-use-efficiency, but not necessarily drought tolerance. Central to the determination of whole-plant water use efficiencies, such as transpiration efficiency (TE, here ratio between aboveground biomass and transpired water; Table 1) or water productivity (WP, here ratio between seed biomass and transpired water; Table 1), is the quantification of water lost by the plant. We have previously shown that leaf-level WUE is not representative of absolute vegetative (rosette) water use (VWU), or biomass production (Ferguson *et al*. 2018), as the transpiring leaf surface is a major upscaling factor. We also demonstrated in a few selected ecotypes and mutants that differences in life-time plant water use (PWU, Table 1) and plant-level water-use efficiency (TE and WP) exist (Bechtold *et al*. 2010, 2013). This is an important consideration since water use is maintained for an extended period following floral transitioning (Bechtold *et al*. 2013), however little is known about the underlying molecular mechanisms of the variation in PWU and TE/WP. In Arabidopsis, the measurement of life-time PWU has received little attention, mainly due to the difficult and time consuming nature of manually phenotyping PWU on a daily basis for the majority of the lifetime of the plant (Bechtold *et al*. 2010, 2013). As plants begin to develop stalks and flowers, automated watering systems (Granier & Tardieu 2009; Tisné *et al*. 2013) would cause considerable disturbance of the tall structures. Conversely, non-conveyor belt platforms (Halperin, Gebremedhin, Wallach & Moshelion 2017), or a manual approach involving careful handling of flowering plants limits the potential for harmful effects occurring due to movement and touch induced changes (Van Aken *et al*. 2016). From limited studies of this nature, the C24 ecotype has emerged as drought tolerant and highly water use efficient (Bechtold *et al*. 2010), additionally it demonstrates resistance to numerous abiotic and biotic perturbations (Brosché *et al*. 2010; Lapin *et al*. 2012; Xu *et al*. 2015; Bechtold *et al*. 2018).

Our recent study of 35 Arabidopsis ecotypes confirmed the above-described uniqueness of C24 in uniting several desirable water use and drought response traits (Ferguson *et al*. 2018). To build upon these findings, we set out to ascertain whether PWU of C24 was reduced compared to other ecotypes and whether this had a heritable and genetically discernible basis. We therefore employed a C24 x Col-0 recombinant inbred line (RIL) population (Törjék *et al*. 2006) to identify QTLs that underlie the natural variation of these traits. However, due to the difficulties of manually phenotyping PWU, development of a suitable proxy trait was required to phenotype the mapping population in a high-throughput manner. Arabidopsis represents an ideal system through which to develop and evaluate the usefulness of proxy traits, such as *WUEi*, δ^13^C, flowering time, VWU and biomass parameters for predicting PWU and whole-plant water-use efficiencies. To this end, we assessed the usefulness of this suite of traits for acting as proxies to predict whole-plant water-use efficiencies (TE and WP, see Table 1) in a set of 12 summer annual ecotypes. A highly accurate proxy trait was subsequently identified and employed in a forward genetic screen for whole-plant water use traits.

## Materials and methods

### Plant material and plant growth

A selection of 12 facultative summer annual *Arabidopsis thaliana* (Arabidopsis) ecotypes (Table S1) and 164 RILs derived from a cross between ecotypes Col-0 and C24 (Törjék *et al*., 2006) were employed to assess the natural variation of long-term plant water use. The genetic map and genotype information for the RIL population are as described in Törjék *et al*., 2006 (Table S2). The Col-0 x C24 RIL mapping population was used to identify QTL relating to key traits associated with water use. Detected QTL regions of interest were further investigated using near-isogenic lines (NILs) that captured Col-0 alleles in a homogenous C24 genomic background and vice versa (Törjék *et al*. 2008). The ecotypes, RILs, and NILs were phenotyped for water-use (VWU and PWU), flowering time, and above ground biomass parameters. Additionally, the 12 ecotypes and NILs were phenotyped for δ^13^C (Fig. 1). Plants were sown in peat-based compost (Levington F2+S, The Scotts Company, Ipswich, UK.) and stratified at 4°C in darkness for 4 days. After stratification plants were grown in a growth chamber at 23°C under short-day (8h:16h; light:dark) conditions, under a photosynthetically active photon flux density (PPFD) of 150 ± 20 μmol m^-2^ s^-1^, and at 65% RH (VPD of 1kPa, Fig. 1). Plants were transferred to the glasshouse at distinct stages depending on the applied watering regime (see below and Fig. 1). Within the glasshouse, the environmental conditions were variable, as temperature and external light cycles fluctuated during the experimental periods. Supplemental lighting was maintained at a minimum PPFD threshold of ~200 *μ*mol m^-2^s^-1^ at plant level for a 12h day (long day conditions). Plants were watered according to the different watering regimes (see Fig. 1), and their positions within the two growth environments (short- and long day) were changed daily. In this study, we deliberately opted for transitions between short- and long-day conditions (growth chamber to glasshouse) without a vernalization period, which resulted in delayed flowering compared to some studies. This decision was taken as physiological measurements (snapshot measurements for *WUEi*) required a minimal rosette size that would normally not be achieved in vernalized plants.

**Figure 1.**
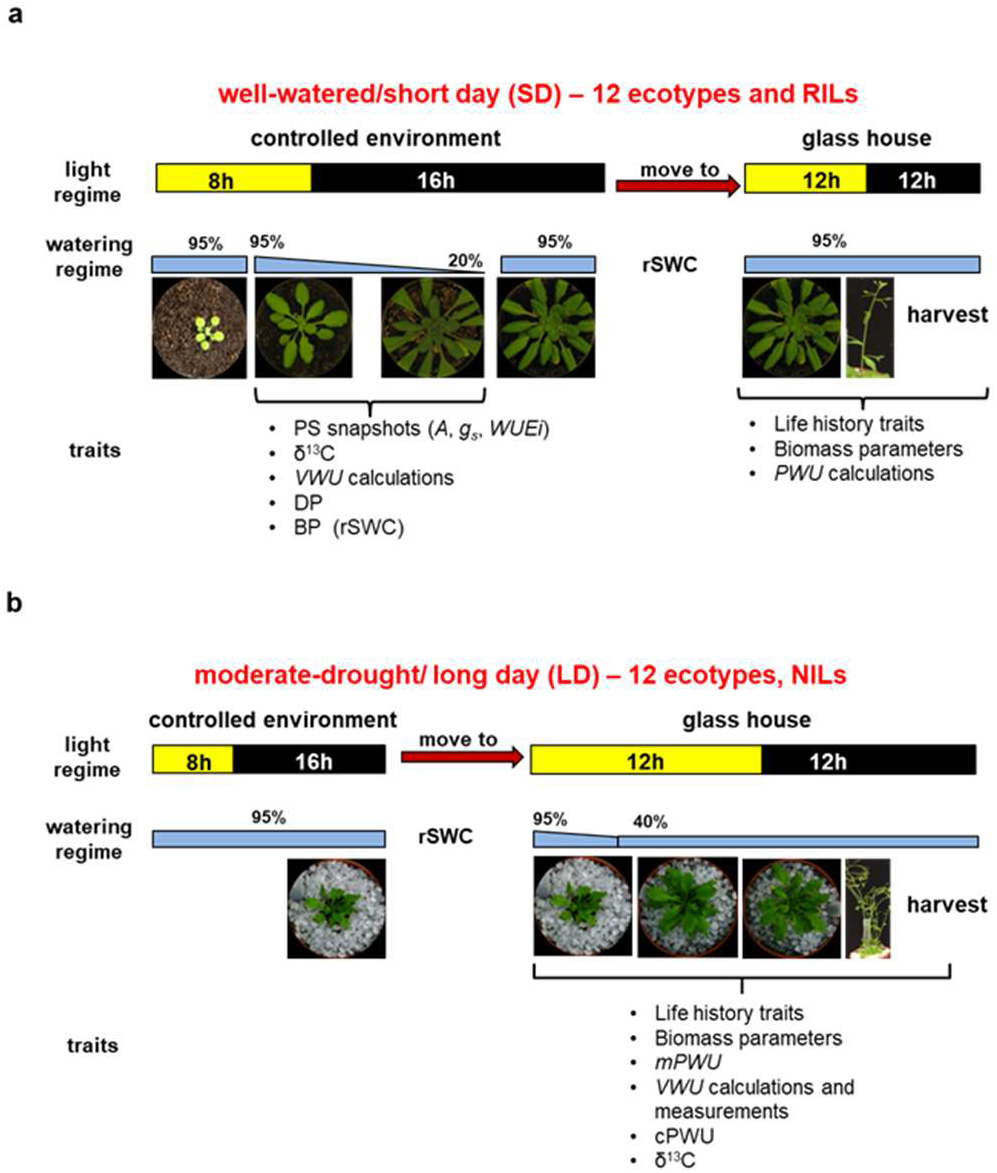
Overview of growth conditions and watering experiments. **a** - short dehydration experiment carried out on 12 ecotypes and the RIL population. Plants were grown for most of their lifespan under short-day (65 days) and well-watered conditions with a short dehydration period to assess plant water use and drought sensitivity, and **b** - continuous maintenance of moderate drought experiment carried out on 12 ecotypes and NILs. Plants were grown for most of their lifespan under long-day and moderate drought conditions (40% rSWC, Bechtold *et al* 2010, 2013). VWU – vegetative water use, PWU – life-time plant water-use, DP-dehydration plasticity. See Table 1 for glossary of terms.

#### Watering regimes

##### (i) Short term dehydration experiment for the determination of vegetative water use (VWU)

All lines undergoing a short-dehydration experiment were grown in the growth chamber in 6cm diameter (0.11L) pots for the determination of vegetative rosette water-use (VWU) as described in (Ferguson *et al*. 2018). Briefly, at 50 days post sowing, plants were left to progressively dry to 20% relative soil water content (rSWC), at which point they were re-watered and transferred from the controlled environment room to the glasshouse for flowering time determination and seed production. VWU was calculated as the slope of the linear regression of the rate of drying from 95 – 20% *rSWC* (lasting between 10 – 12 days; Fig. 1a, Table 1). Plants were transferred to the glasshouse after re-watering and maintained well-watered to determine flowering time and the number of rosette leaves at bud initiation. Plant biomass components were separated and measured as rosette biomass (vegetative biomass), chaff biomass (stalks and pods; reproductive biomass), and seed yield (reproductive biomass), and the sum of all biomass components produced the total above ground biomass value. PWU was calculated as VWU multiplied by the time it took from germination to flowering to generate calculated lifetime plant water use (cPWU, Table 1). WP was calculated as seed biomass divided by either calculated or measured life-time water use (cWP or mWP, Table 1). This watering regime is designated as short day (SD), as plants spend most of their life time under short day conditions (~65 days)

##### (ii) Continuous maintenance of moderate drought for determination of life-term plant water use (PWU)

For the determination of PWU, 8-cm diameter (0.3L) pots were filled with the same volume of soil following the experimental setup as described in Bechtold *et al*. (2010). The soil surface was covered with 0.4 cm diameter polypropylene granules to limit soil evapotranspiration. Plants were germinated in the previously described growth chamber before being transplanted into individual pots 12 days after sowing at the initiation of the rosette growth stage (Boyes *et al*. 2001). Four days after being transferred into individual pots, plants were moved into the glasshouse, where pots were weighed daily (Kern PCB, 350-3 balance) to determine and maintain the pots at a moderate drought level of 40% rSWC (Bechtold *et al*. 2010). Daily water use was recorded after plants were transferred into the glass house. Control pots without plants were also measured daily to estimate evaporation from the soil surface. Flowering time and number of leaves at bud initiation were recorded, and once the final flower had opened, watering ceased, and plants were bagged for harvesting. During harvest the vegetative (rosette) and reproductive (stalks, pods, and seeds) biomass components were separated. mPWU was determined as the sum of water added every day until bagging, minus the water lost through evaporation from control pots. This parameter is also termed measured plant water use (mPWU) in order to distinguish it from cPWU (Table 1). This watering regime is denoted as long-day (LD), as plants only spend 16 days from germination under short day conditions, the remaining time plants were grown under long day conditions (Fig. 1b).

### Estimating drought sensitivity (DS)

For analysing in more detail the data used for calculating VWU, we applied the Davies test (Davies, 2002) and segmented regression analysis as part of the segmented package in R (Muggeo 2017) in order to test (i) for a significant difference in slope parameter and (ii) for the breakpoint in the regression. This analysis produced the breakpoint in the drying period and the slopes before (stage 1) and after (stage 2) the breakpoint. VWU plasticity was calculated as the slope before the breakpoint (stage1; supposed to represent transpiration under control conditions) - slope after breakpoint (stage2; supposed to represent transpiration under drought conditions) / slope before breakpoint (stage1). Both breakpoint (in terms of rSWC) and VWU plasticity were used to estimate the drought sensitivity as per Ferguson *et al*. (2018).

### Physiological measurements

*(i) Photosynthetic rate (snapshot measurements) in the short dehydration experiment* Instantaneous measurements of net CO_2_ assimilation rate (A), and stomatal conductance to water vapour (*gs*) and transpiration rate (E) were taken on leaf 7, using an open gas exchange system (PP Systems, Amesbury, MA, USA). Leaves were placed in the cuvette at ambient CO_2_ concentration (C_*a*_) of 400 μmol mol^-1^, leaf temperature was maintained at 22 ± 2 °C and vapour pressure deficit was ca. 1 kPa and irradiance was set to growth conditions (150 μmol m^-2^ s^-1^). A reading was recorded after the IRGA conditions had stabilized (ca. 1.5 min), but before the leaf responded to the new environment (Parsons, Weyers, Lawson & Godber 1997). Instantaneous water use efficiency *WUEi* was estimated as A/E.

#### (i) Delta carbon 13 analysis

The carbon isotope composition (δ^13^C) of bulk leaf material was assessed for the 12 ecotypes comprising the SD experiment (well-watered samples), and the NILs and parental lines from the continuous moderate drought experiment. The harvested leaves had developed during moderate drought stress (40% rSWC). δ^13^C was measured as described in Roussel *et al*. (2009) and Ferguson *et al*. (2018). δ^13^C was calculated as: (R_s_ – R_b_)/R_b_ x 1000, where R_s_ and R_b_ represent the ^13^C/^12^C ratio in the samples and in the Vienna Pee Dee Belemnite standard respectively (Craig, 1957).

### Statistical Analysis

All statistical analyses were performed within the R software environment for statistical computing and graphics (R Core Team 2015). Experiments using the RIL population were performed across several blocks over a period of two years. Each temporally divided block contained the two parental ecotypes and between 20-40 RILs. One-way analysis of variance (ANOVA) comparison of means tests were performed across all lines and all blocks to determine the existence of experimental block effects that could potentially confound further analysis and the QTL mapping. Best linear unbiased predictors (BLUPs) were extracted using the following general linear mixed model (GLMM): Y = E + B + Residual (Error) variance, where Y represents the phenotypic trait parameter of interest, and both E (Ecotype) and B (Experimental block) are treated as random effects, while controlling for fixed effects, i.e. temporal block effects (Lynch & Walsh 1998). Predicted means were obtained for each trait and for each RIL by adding the appropriate BLUP value to the population mean. Predicted means were employed for all subsequent analyses involving the RILs and for QTL mapping. The GLMMs allowed for the determination of phenotypic (V_P_) and genotypic (V_G_) variation for all trait parameters. These parameters were used to obtain estimates of broad sense heritability (*H^2^*) as V_G_/V_P_.

### QTL Mapping

We mapped for QTLs underlying all assessed parameters using the qtl R package (Broman *et al*. 2003; Broman 2009). The Lander-Green algorithm (Lander & Green 1991) i.e. the hidden Markov model technology, was used to re-estimate the genetic map using the Kosambi map function to convert genetic distance into recombination fractions with an assumed genotyping error rate of 0.0001. The re-estimated genetic map, based on the lines incorporated in this study, was preferred to the original genetic map, which was based on over 400 RILs. The hidden Markov model technology and Kosambi map function were further employed to calculate the probabilities of true underlying genotypes at pseudo-marker points between actual markers based on observed multipoint marker data, whilst allowing for the same rate of genotyping errors. Genotypes were calculated at a maximum distance of 2 cM between positions.

Multiple QTL mapping was performed using the predicted means derived from BLUPs. The best multiple QTL models were identified via the Haley-Knott regression method (Haley & Knott 1992) using genotype probabilities at both genetic markers and calculated pseudo-markers. We used the Haley-Knott regression method because the genetic map marker density was relatively high (Average inter-marker distance: 3.87 cM). We additionally tested the expectation-maximization and multiple-imputation methods for QTL mapping but did not notice a perceptible difference in QTL effect size or position between these three methods (Fig. S4).

10,000 permutations were used to determine LOD significant thresholds for incorporating both additive QTL and epistatic interactions at an experiment-wise α = 0.05. Automated stepwise model selection was performed, where additive and epistatic QTL were scanned for at each step (Manichaikul, Moon, Sen, Yandell & Broman 2009). The penalties for the stepwise model selection were derived from a two-dimensional genome scan. Finally, the positions of detected QTLs were refined, and the model was fitted with ANOVA to calculate the effect size, percentage variance explained, LOD score for each QTL, and penalized LOD score for each model. Interval estimates of all detected QTLs were obtained as 95% Bayesian credible intervals.

Following MQM, the log_10_ ratio comparing the full model and the single QTL model from the two-dimensional genome scan was directly assessed to test for the presence of an epistatic interaction between the two main effect QTL for cPWU. To assess the impact of flowering time, vegetative biomass, and VWU on the QTL mapping for cPWU, we performed single-QTL mapping for cPWU using these traits as individual covariates.

### *Genotyping using* InDel *markers*

Insertion-deletion markers (InDels) polymorphic between Col-0 and C24 alleles of FRI and FLC were obtained to address the hypothesis that these genes underlie the two major QTLs detected. A 16-bp deletion in the Col-0 allele of FRI was scored using primers developed by Johanson *et al*. (2000). A 30-bp deletion in the Col-0 allele of *FLC* was scored using primers developed by Gazzani *et al*. (2003). InDel markers with a single PCR band for both InDels (Fig. S1a; Table S3) were assayed by qPCR and high-resolution melting (HRM) genotyping using the CFX96^TM^ Touch Real-Time PCR Detection System (BIO-RAD). This information for 138 individuals of the RIL population and both parents was subsequently integrated into the re-estimated genetic map (Fig. S1b, Table S4).

### Analysis of publicly available RNAseq and microarray datasets

Publicly available RNAseq (Xu *et al*. 2015; GSE61542) and microarray datasets of C24 and Col-0 (Bechtold *et al*. 2010, E-MEXP-2732) were analysed for differentially expressed genes. These datasets were compared with the protein coding genes within mapping intervals using VENNY (Oliveros 2007).

### RNA extraction and gene expression analysis by qPCR

Leaves of a minimum of four biological replicates were harvested from the NILs and both parental lines at 26 and 43 days post germination, and frozen in liquid nitrogen. Total RNA was extracted using Tri-reagent (SIGMA, Aldrich, UK) according to the manufacturer’s instructions. For cDNA synthesis, 1 μg of total RNA was treated with RNase-free DNase (Ambion) according to manufacturer’s instructions and reverse transcribed as previously described (Bechtold *et al*. 2008). Quantitative real-time PCR (qRT-PCR) was performed using a cybergreen fluorescence based assay as described previously (Bechtold *et al*. 2008). Gene-specific cDNA amounts were calculated from threshold cycle (Ct) values and expressed relative to controls and normalized with respect to Actin and Cyclophilin cDNA according to Gruber, Falkner, Dorner & Hämmerle (2001). To calculate the standard error of the calculated ratios of fold differences for gene expression data, the errors of individual means were combined "in quadrature", and the final ratio was a combination of the error of the two-different means of the NILs and Col-0 samples. The primers used for RT-qPCR can be found in Table S3.

## Results

We used a selection of 12 facultative summer annual ecotypes of Arabidopsis that previously demonstrated variation for drought sensitivity and water use associated traits (Table S1, Ferguson *et al*. 2018), as well as a RIL mapping population and associated NILs (BC4F3-4) to examine natural variation of PWU and above ground biomass allocation (Table S2, Table S5). The assessment of natural variation for VWU, PWU, biomass accumulation and drought sensitivity was followed by QTL mapping to establish the genetic basis of these traits. Two experimental setups were used as part of this study: (i) **12 ecotypes and RILs** - a short dehydration experiment under predominantly short-day conditions to measure a range of leaf level *WUE* parameters (*WUEi*, δ^13^C), VWU, flowering time, biomass parameters and drought sensitivity (Fig. 1a, Ferguson *et al*. 2018), and (ii) **12 ecotypes and NILs** - a continuous moderate drought experiment under predominantly long-day conditions, during which *rSWC* was maintained at moderate drought levels (~40% *rSWC*) to measure leaf level *WUE* parameters (δ^13^C), VWU, PWU, flowering time and biomass parameters (Bechtold *et al*. 2010; Fig. 1b).

### Identification of a proxy trait for lifetime (plant) water use (PWU)

We analysed a range of parameters associated with plant water status by performing a short dehydration as well as a continuous maintenance of moderate drought experiment on 12 selected Arabidopsis ecotypes (Fig. 1, Table 1). We determined VWU (Ferguson *et al*. 2018; Fig. 1a, Table 1), lifetime PWU (Fig. 1b, Table 1), flowering time, above ground biomass parameters, δ^13^C, and calculated whole-plant water-use efficiency parameters, namely TE and WP (Table1; Fig. 1a, b; Bechtold *et al*. 2010, 2013, 2016; Ferguson *et al*. 2017). Both δ^13^C and *WUEi* measurements were taken to determine the influence of leaf-level processes on whole plant traits (i.e. transpiring leaf surface area), however we did not observe a significant relationship with whole-plant water-use efficiency parameters such as TE and WP (Fig. S2). We continued to focus on the determination of lifetime PWU and the genetic dissection of PWU and productivity traits, instead of the leaf-level *WUE* parameters, δ^13^C and *WUEi*.

Our usual approach of a manual determination of PWU (Fig. 1b) requires the weighing and watering of individual pots until the terminal flower has opened (Bechtold *et al*. 2010). The manual determination of PWU is challenging and time consuming (see Introduction), thus to facilitate large-scale manual screening of PWU of the mapping population, we first set out to identify an adequate proxy. We compared biomass production, flowering time, VWU and PWU between the short-dehydration and continuous moderate drought experiment carried out on the 12 Arabidopsis ecotypes (Fig. 1). The continuous moderate drought experiment revealed that measured PWU (mPWU) was significantly correlated with both flowering (Fig. 2a) and vegetative (rosette) biomass (Fig. 2b, Table S6). Based on these relationships we developed the proxy parameter “*calculated life time (plant) water-use* (cPWU)”, as a product of VWU and flowering time:

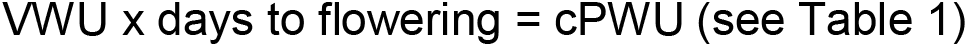

**Figure 2.**
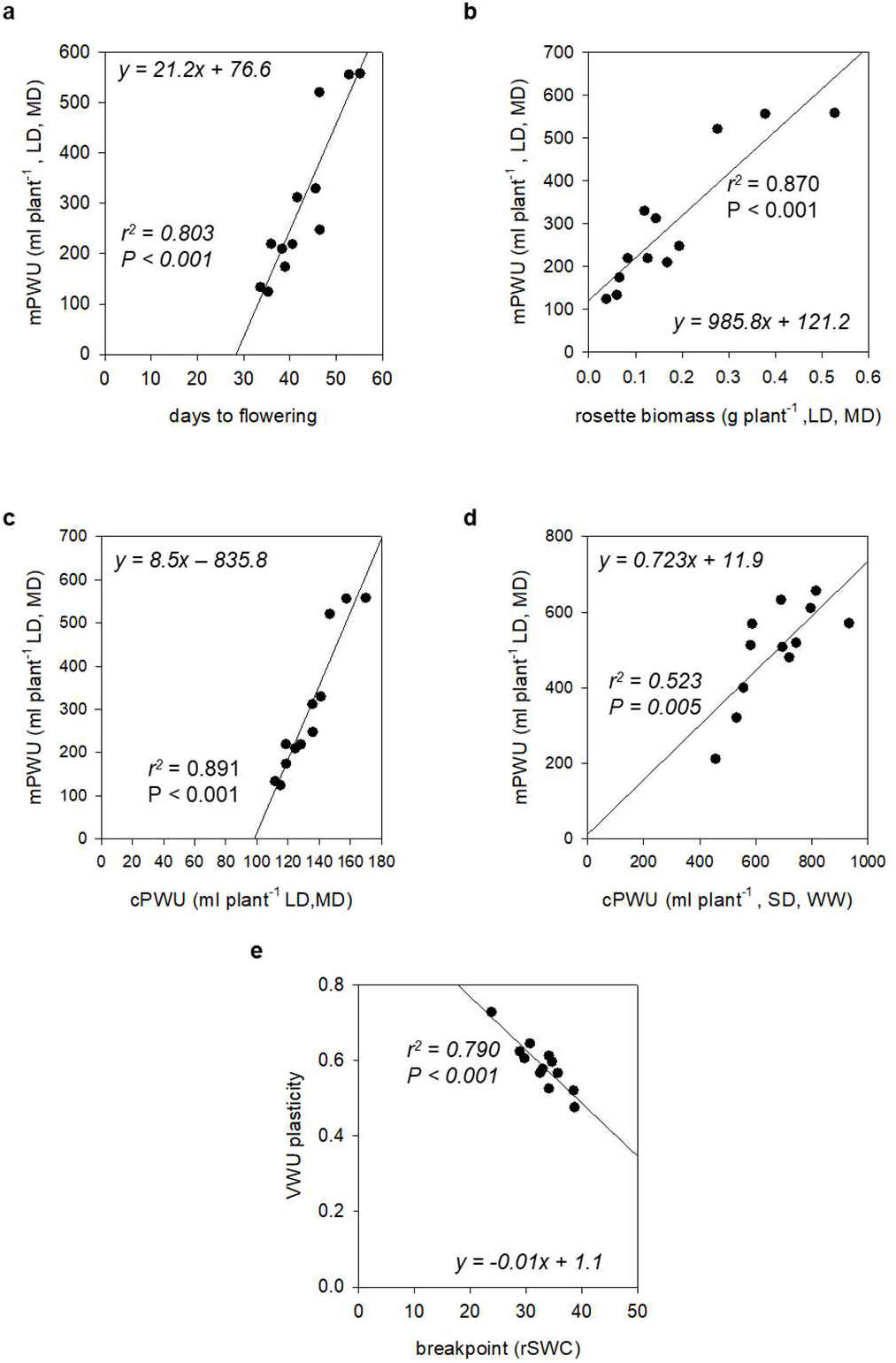
Lifetime water-consumption and performance parameters in 12 selected ecotypes. **a** - relationship between days to flowering and mPWU, **b** – relationship between vegetative biomass and mPWU, **c** – relationship between cPWU and mPWU within the same experiment, **d** - relationship between cPWU and mPWU between two independent experiments: long-day, moderate-drought (LD, MD), and short-day, well-watered (SD, WW). The lines represent the equation of the linear regression model, and **e** - relationship between the breakpoint in dehydration response and VWU plasticity. The P-value of the slope parameter and adjusted *r*^2^ value associated with the linear model are provided for each association.

The continuous moderate drought experiment allowed us to directly relate mPWU with cPWU, which showed a highly significant positive correlation within the experiment (Fig. 2c). In addition, the correlation between mPWU with cPWU was tighter than the correlations with rosette biomass and flowering time (Fig. 2a, b). Importantly, a significant correlation between calculated and measured PWU was also observed when comparing mPWU from the continuous moderate drought experiment under long-day conditions, with cPWU of a short-dehydration experiment under short-day conditions (Fig. 2d). Therefore, we reasoned that PWU calculated from flowering time in a short-dehydration experiment would provide a robust estimate of mPWU.

Furthermore, the short-dehydration approach allowed us to quantify the drought responses of individual ecotypes by calculating the threshold at which plants enter drought stress (breakpoint) and the plasticity of the drought response (VWU plasticity; Ferguson *et al*. 2018). The breakpoint negatively correlated with the VWU plasticity, indicating that lines responding to drought stress at higher rSWC, showed less absolute change in transpiration throughout the dehydration period, and therefore exhibited reduced VWU plasticity (Fig. 2e). Therefore, a short dehydration experiment allowed us to not only screen and dissect the genetic basis for the natural variation of cPWU and biomass, but also assess drought response parameters at the same time.

### The genetic dissection of cPWU, drought response and biomass parameters

Short dehydration experiments (Fig. 1a) were subsequently performed on 163 individuals of the Col-0 x C24 RIL population (Table S2) including both parents. To control for experimental block effects, BLUPs were extracted and predicted means were calculated for all traits. The variation in predicted means for all traits was not significantly different from what would be expected of a normal distribution (P > 0.05; Kolmogorov-Smirnov normality test) and all traits demonstrated transgressive segregation (Fig. S3). We calculated genetic variance (V_G_), total phenotypic variance (V_P_) and broad sense heritability (*H^2^*), where all 13 traits assessed demonstrated variation that had a significant heritable basis within the RIL population (Table 2).

**Table 2.**
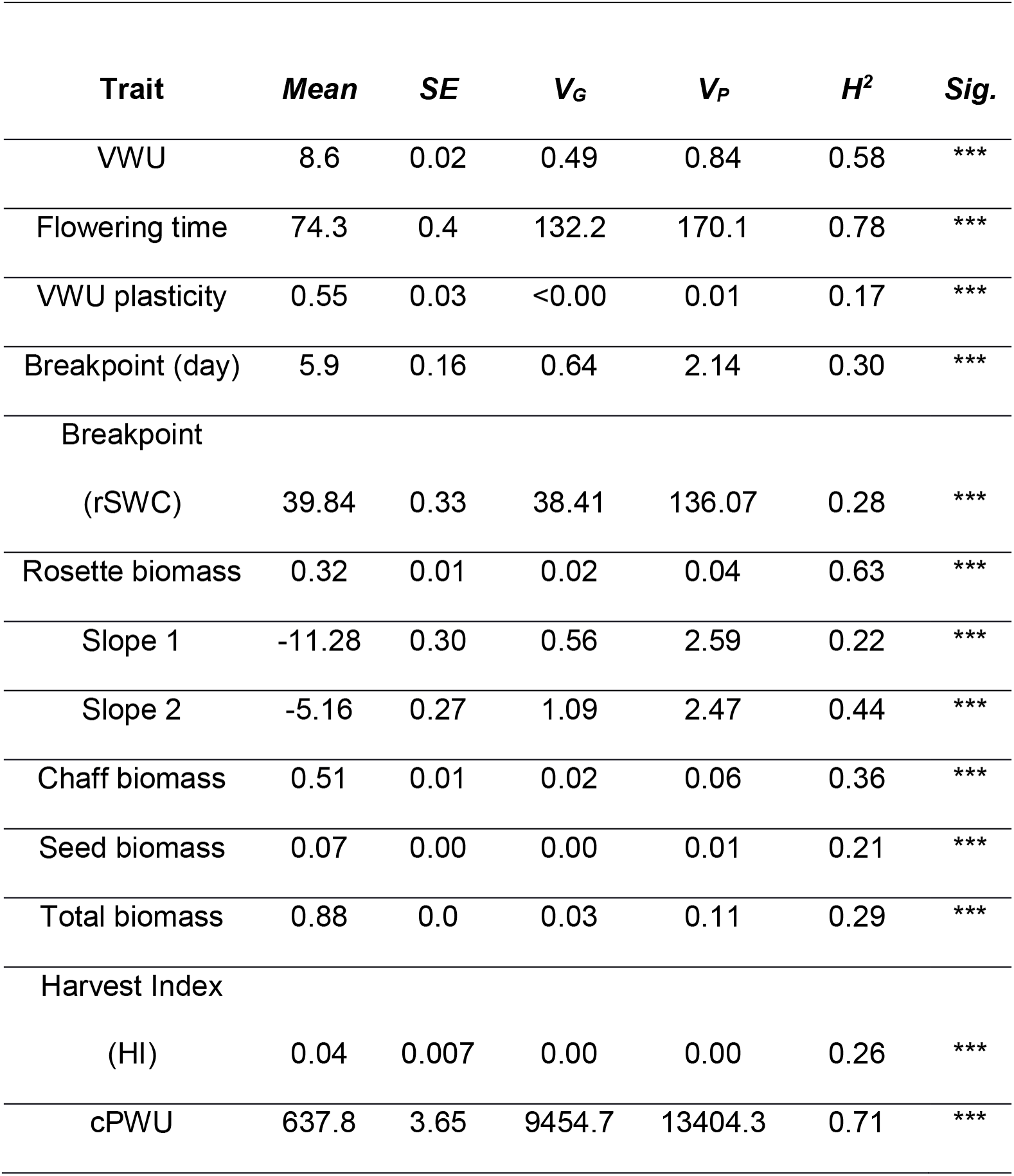
Genotypic and phenotypic variation of the twelve traits assessed as part of the QTL mapping. The true (arithmetic) mean, standard error (SE), genetic variance (VG), phenotypic variance (VP), broad sense heritability (*H^2^*), and significance of *H^2^* (Sig.) are provided for all traits. n.s – not significant, *** indicates significant heritability at the p < 0.001 level.

Adjusted linkage maps were constructed based on the individuals used for mapping. Analyses indicated that 97.5% of the markers had been genotyped for all the RILs, and we observed a virtually even split in the allelic form of these markers, with 50.3% coming from the Col-0 parental line and 49.7% from the C24 parental line. To identify the genetic variation that causes the observed phenotypic variation in VWU, cPWU, flowering time, productivity and drought sensitivity traits, multiple QTL mapping was performed (MQM; see Material and methods) on a minimum of 163 selected individuals. No significant QTL models were identified for seed biomass (Fig. S5a), dehydration response (VWU plasticity; Fig. S5b), and the breakpoint (Fig. S5c). For VWU, FT, cPWU and slope 1, a total of 9 main effect QTLs were detected (Fig. 3; Table 3; p <0.001). The percentage of phenotypic variance explained for the cPWU QTLs ranged from 4.8 to 25.5%, for flowering time from 3.6 to 18.2 %, and for VWU from 3 and 5% (Table 3). Consistent with the previously observed significant correlation between flowering time and mPWU (Fig. 2a), there was co-localization between the two main effect QTLs on chromosomes 4 and 5 for flowering time and cPWU (Fig. 3a, b; Table 3). The strong positive correlation observed between flowering time and PWU suggests that the co-localizing QTLs for these traits were likely to represent the same genes or linkage between causal genes. On the other hand, QTLs detected for VWU did not co-localise with flowering time QTLs (Table 3, Fig. 3).

**Figure 3.**
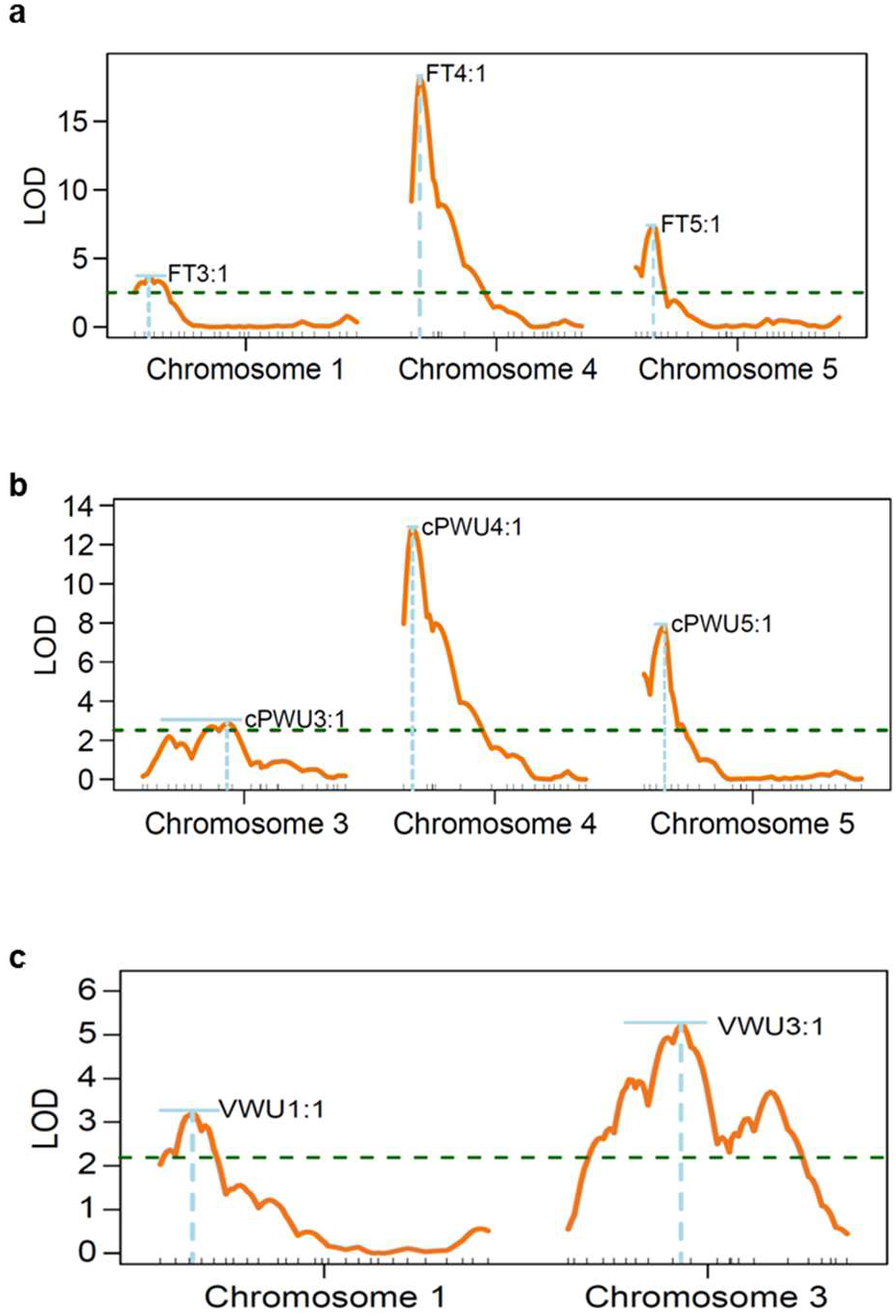
QTL mapping. LOD profiles for whole chromosomes were significant QTL are located according to multiple QTL mapping (MQM). **a** - LOD profiles for three significant QTLs underlying variation for flowering time (FT), **b** - LOD profiles for three significant QTLs underlying variation for calculated lifetime plant water-use (cPWU), and c - LOD profiles for vegetative water use (VWU).

**Table 3:** Locations and effect sizes for the significant QTL arising from the QTL mapping via a MQM for water use, harvest index and flowering time. The QTL names are given as the trait followed by the chromosome location. The position in cM, LOD score (LOD), proportion of total genetic variation, 95% Bayesian credible interval, P-value, and additive genetic effect provided for all significant QTLs.

| QTL | Position (cM) | LOD score | Proportion of total genetic variation | 95% Bayesian credible interval (cM) | P-value | Additive genetic effect (SE) |
| --- | --- | --- | --- | --- | --- | --- |
| VWU:1 | 10 | 3.210 | 7.561 | 0-18 | < 0.000 | 0.19 (0.05) |
| VWU:3 | 35 | 5.215 | 12.643 | 18-43 | < 0.000 | -0.25 (0.05) |
| FT:1 | 7 | 3.578 | 5.346 | 1-14 | < 0.000 | -2.86 (0.69) |
| FT:4 | 3.69 | 18.177 | 34.769 | 3-5 | < 0.000 | 6.75 (0.64) |
| FT:5 | 8 | 7.334 | 11.658 | 5-9 | < 0.000 | -3.95 (0.64) |
| cPWU:3 | 36 | 2.851 | 4.799 | 8-42 | < 0.000 | -21.99 (6.01) |
| cPWU:4 | 4 | 12.785 | 25.455 | 2-6 | < 0.000 | 48.79 (5.79) |
| cPWU:5 | 8.82 | 7.811 | 14.283 | 5.25-10 | < 0.000 | -35.90 (5.69) |
| Slope1:1 | 32.61 | 2.511 | 6.889 | 0-86.829 | <0.000 | 0.16 (0.05) |

When performing single QTL-mapping for cPWU whilst incorporating flowering time as a covariate in the analyses, the main effect QTL on chromosomes 4 and 5 are not detected, however the QTL on chromosome 3 that is also detected when mapping for VWU becomes more significant (Fig S6c). Similarly, when incorporating vegetative biomass as a covariate, the effect of these QTL is reduced, however they are still significant (Fig S6b). Incorporating VWU as covariate, removes the importance of the QTL on chromosomes 3 and heightens the significance of the QTLs on chromosomes 4 and 5 (Fig. S6d).

The two significant cPWU and flowering time QTLs on chromosomes four and five (Fig. 3a, b) contained two well characterised flowering time genes, *FRIGIDA* (*FRI*, Chromosome 4; AT4G00650) and *FLOWERING LOCUS C* (*FLC*, Chromosome 5; AT5G10140). The ecotype Col-0 possesses a non-functional allele of *FRI* (*fri*) and a functional allele of *FLC* (*FLC*), while the ecotype C24 contains a functional allele of *FRI* (*FRI*) and a weak allele of *FLC* (*flc*; (Johanson *et al*. 2000; Michaels, He, Scortecci & Amasino 2003). A significant epistatic interaction was detected between these QTLs when comparing the full model to the model which just accounts for one of these two QTL (Fig. S7). Transcriptional levels of *FLC* are positively regulated by *FRI* (Deng *et al*. 2011), thus the epistatic interaction between these QTL further suggests that *FRI* and *FLC* are the causal genes. InDel markers were designed for both candidate genes and the RIL population was scored for the allelic variant of both genes (see Materials and Methods). This information was incorporated into the genotypic data and the genetic map was re-estimated, which demonstrated that *FRI* and *FLC* were present between the markers that flanked the main effect QTLs on chromosomes four and five respectively (Fig. S1b). The RIL population was sub-divided according to the different allelic combination of *FRI* and *FLC* of each individual line (Table S4) to confirm the importance of the functionality of these genes on the traits of interest here.

### The genetic action of non-functional and weak alleles of *FRI* and *FLC* reduces water-use

We determined the allelic state of *FRI* and *FLC* in all RILs and divided the population into four groups: i) *fri*: *FLC* (Col-0), ii) *FRI*: *FLC*, iii) *fri*: *flc*, and iv) *FRI*: *flc* (C24). One-way ANOVA comparisons of means and post-hoc Tukey tests were performed to determine the effect of different allelic combinations on water use and plant development (Fig. 4). There were significant and parallel differences in cPWU and flowering time between the four groups (Fig. 4a, b). Possessing non-functional and weak alleles of *FRI* and *FLC*, respectively, significantly reduced flowering time and cPWU (Fig. 4a, b).

**Figure 4.**
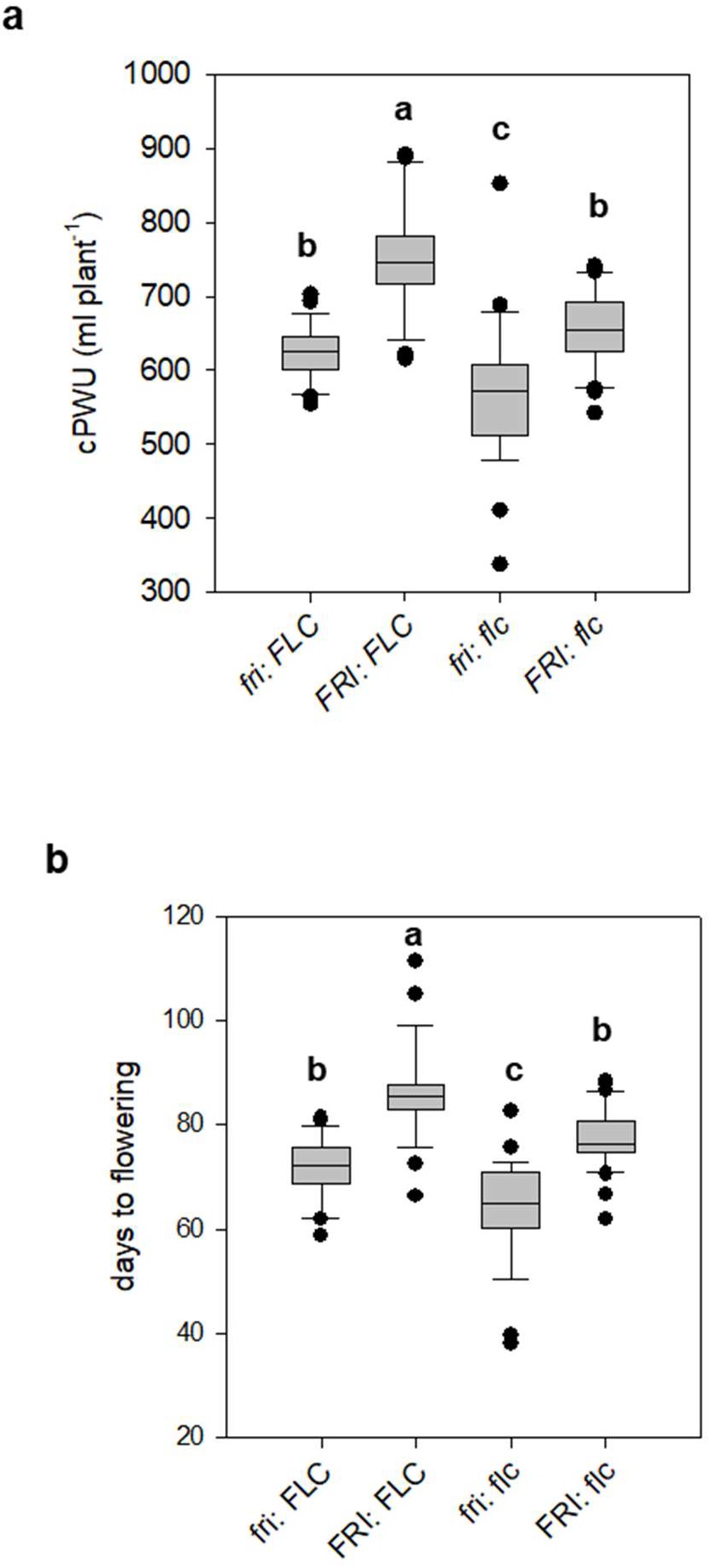
Trait performances of genotypes harbouring different allelic combinations of the FRIGIDA and FLOWERING LOCUS C genes in RILs. Boxplots describing the variation for traits assessed for the 4 groups based on allelic combination of both *FRI* and *FLC*, a – cPWU, **b** - days to flowering. The letters (a, b, and c) above the boxplot denote the post-hoc Tukey groups, where allelic groups whose letters are different are significantly different from one another for that trait at P < 0.05. The bold line in the centre of the boxplots represents the median, the box edges represent the 25th (lower) and 75th (upper) percentiles, the whiskers extend to the most extreme data points that are no more than 1.5x the length of the upper or lower segment. Outliers are data points that lie outside the 1.5x interquartile range both above the upper quartile and below the lower quartile.

To further test the hypothesis that cPWU is a suitable proxy of mPWU and to confirm that increased life-span through a combination of *FRI* and *FLC* is the main factor underlying PWU, we subsequently obtained NILs that harboured the Col-0 allele of FRI and FLC separately in a homogenous C24 genomic background and vice versa (Table S5). Seven NILs and two parental lines were subjected to a continuous moderate drought experiment, where flowering time, mPWU, VWU, cPWU, productivity parameters, mean daily water use as well as δ^13^C and stomatal conductance were determined (Fig. 1b). The hypotheses regarding cPWU that emerged from the RIL population were essentially confirmed. The combination of both non-functional and weak alleles of *fri* (Col-0) and flc (C24) led to significantly reduced mPWU (Fig S8a), but had no impact on VWU (Fig. S8c,d).

Due to the significant relationship between flowering time and mPWU (Fig. 2a), we assessed whether the different allelic combinations of *FRI* and *FLC* had pleiotropic effects on VWU. There was no significant difference in VWU in both the NILs and RILs under either short-(RILs) or long-day (NILs) conditions (Fig. S8c,d).

Interestingly we observed a significant relationship between mean daily water-use, days to flowering and rosette biomass in the moderate drought experiments for the 12 ecotypes and the NILs (Fig. S9a,b; Fig 5a,b), leading to high mPWU (Fig. S9c; Fig. 5c). Therefore, late flowering ecotypes and NILs appear to sustain increased daily water use over a longer period, which was independent of the allelic combinations of *FRI* and *FLC* (Fig. 5d).

**Figure 5.**
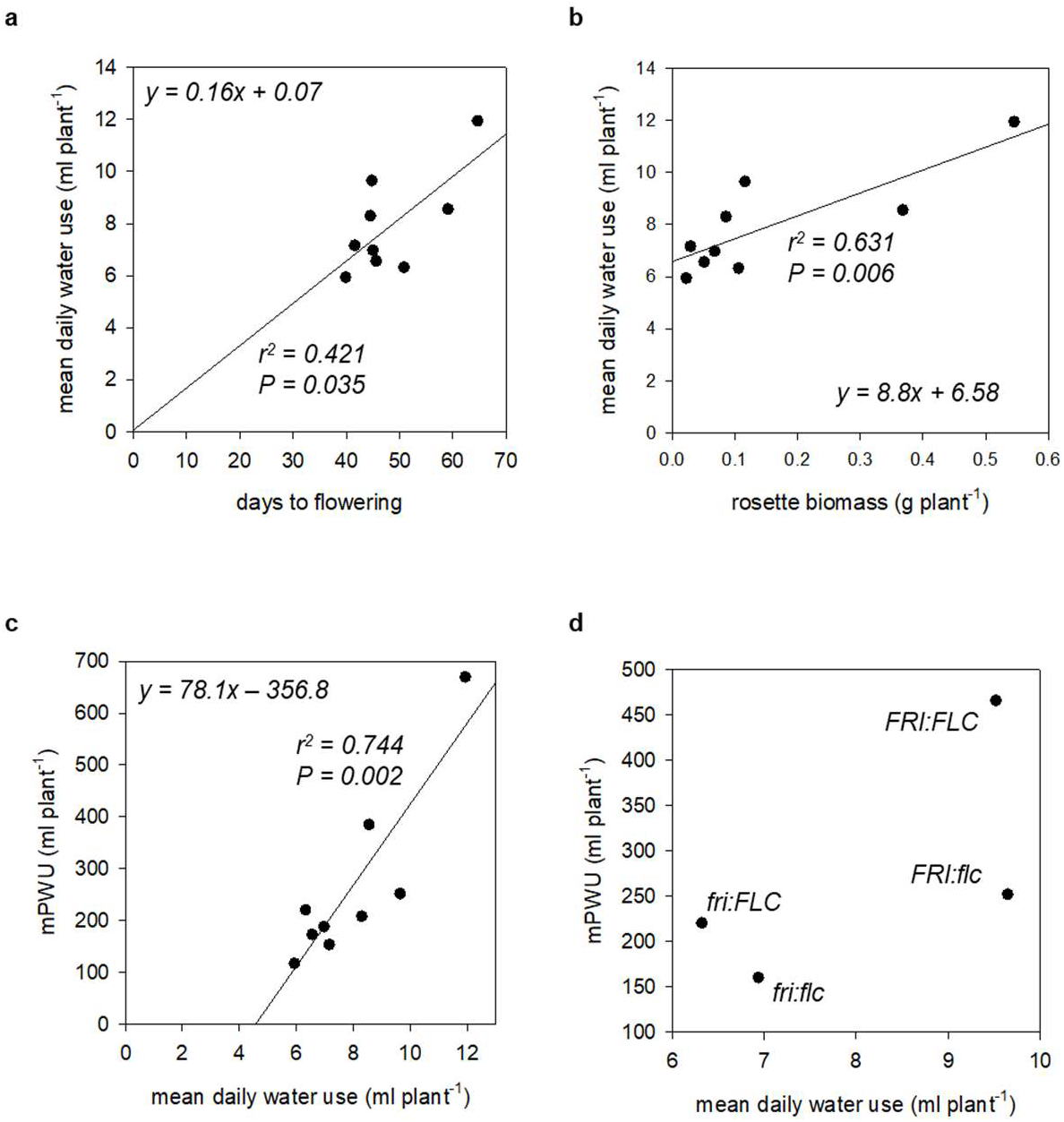
The contribution of mean daily water use in the NILs. **a** - relationship between flowering time and mean daily water use, **b** - relationship between rosette biomass and mean daily water use, **c** - relationship between mean daily water use and mPWU, and **d** - relationship between mean daily water use and mPWU divided into four *FRI*/*FLC* allelic groups tested in the NILs. The linear model of the relationship between mean long-term water use and mean daily water use is provided. R2 and P values are provided where a significant relationship was identified.

δ^13^C, while significantly different between Col-0 and C24, did not show a significant difference among the remaining allelic combinations of *FRI* and *FLC* (Fig. S10a), which suggests that δ^13^C was independent of *FRI* and *FLC*. A significant negative correlation between δ^13^C and stomatal conductance indicated that low g_s_ leads to increased instantaneous WUE (A/gs) (*g_s_*; Fig. S10b; R^2^ = 0.781 p < 0.01), which also coincided with the distinct rosette growth phenotype of C24 (Fig. S10b,d). In addition, the lack of significant QTLs for VWU, VWU plasticity and the breakpoint (Fig. S5) suggests that leaf-level drought responses were not genetically controlled in this mapping population, and therefore independent of the detected genetic control of flowering time. This was confirmed by the non-significant differences in VWU, VWU plasticity and breakpoint for the four allelic *FRI*/*FLC* groups (Fig. S8c,d; Fig. S11a,b).

Importantly, the observation that a combination of *fri* (Col-0) and *flc* (C24) in the NILs led to significantly reduced mPWU (Fig S8a), and significant variation in δ^13^C (Fig S10a) that did not match the variation for mPWU, support our observations from the diverse suite of ecotypes. Taken together, this suggests that cPWU is a reliable proxy for mPWU.

### Biomass variation and distribution is independent of the genetic action of *FRI* and *FLC*, and growth conditions

We also assessed whether the different allelic combinations of *FRI* and *FLC* resulting in significantly different PWU, had pleiotropic impacts on biomass parameters. For example, the decrease in cPWU in the *fri: flc* group did not result in a significant reduction in above ground-, seed- or vegetative biomass in the RILs (Fig. 6a-c) or the NILs (Fig. S12a-c), yet the combination of *FRI:FLC* significantly decreased seed- and increased vegetative biomass (Fig. 6b,c; Fig. S12c). This suggests that the additionally acquired photosynthates acquired by later flowering plants are translocated primarily to vegetative as opposed to reproductive sinks.

**Figure 6.**
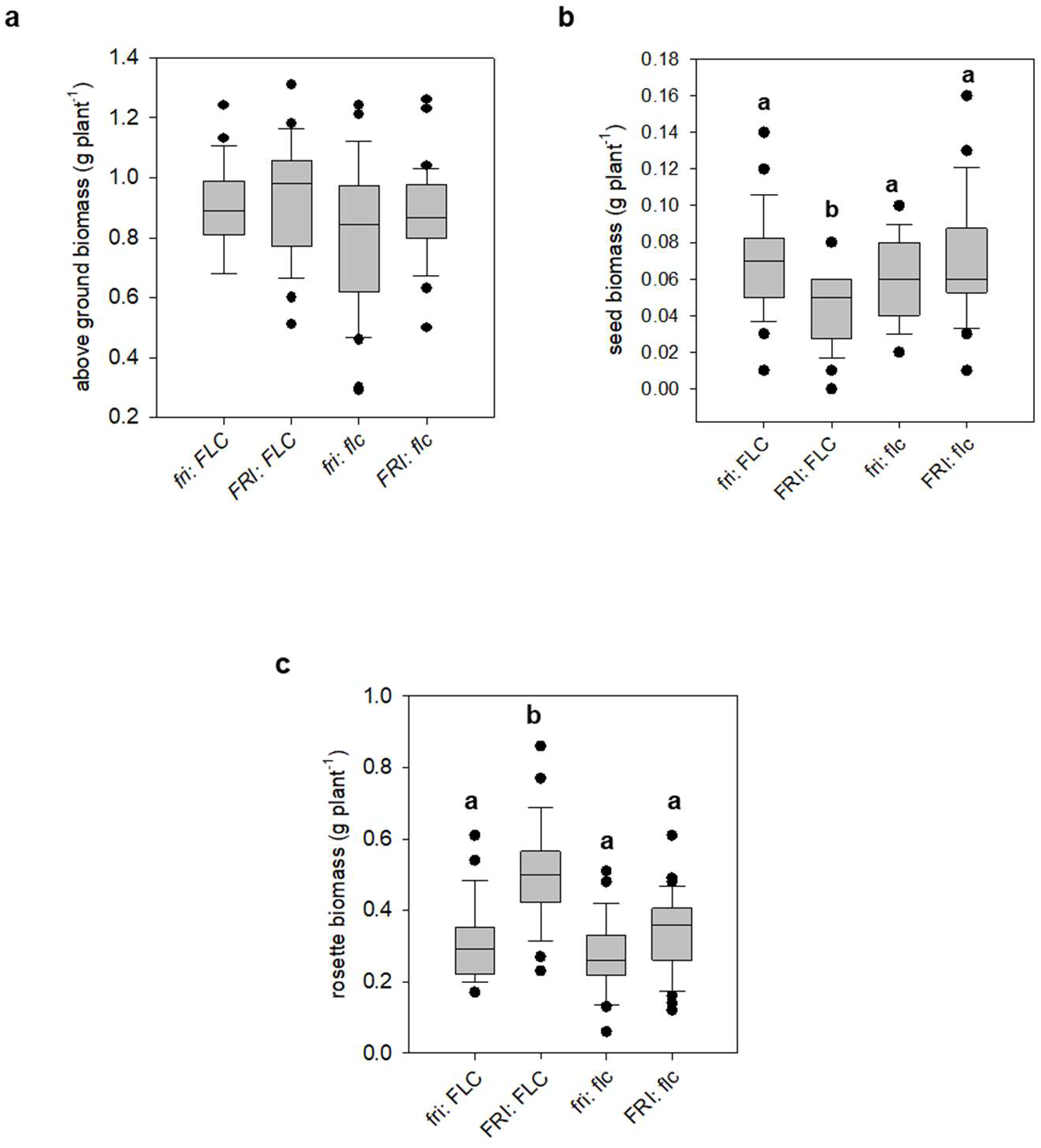
Boxplots of biomass parameters based on allelic combinations of FRI/FLC in the RILs. **a** – above ground biomass, **b** – seed biomass, and **c** – rosette biomass. The letters (a, b, and c) above the boxplot denote the post-hoc Tukey groups, where allelic groups whose letters are different are significantly different from one another for that trait at P < 0.05. The bold line in the centre of the boxplots represents the median, the box edges represent the 25th (lower) and 75th (upper) percentiles, the whiskers extend to the most extreme data points that are no more than 1.5x the length of the upper or lower segment. Outliers are data points that lie outside the 1.5x interquartile range both above the upper quartile and below the lower quartile.

Biomass allocation (HI) showed substantial variation amongst the NIL and the RIL populations (Fig. S13a, b), due to different experimental conditions (short day vs long day, well-watered vs moderate drought). Despite these experimental differences, relative proportions were highly correlated between the well-watered and moderate drought experiments (Fig. 7), suggesting allelic combinations with low HI in the short-dehydration experiments (RILs) also showed low HI in the continuous moderate drought experiment (NILs; Fig. 7a,). Equally, cPWU significantly correlated across the distinct experiments for the different allelic groups (Fig. 7b). A similar relationship for PWU and HI across different experiments was also observed in the 12 natural ecotypes (Fig. 2d, Fig. 7c). This suggests that the distribution of biomass and PWU was independent of environmental growth conditions including watering status and day length in both the mapping population and the natural ecotypes.

**Figure 7.**
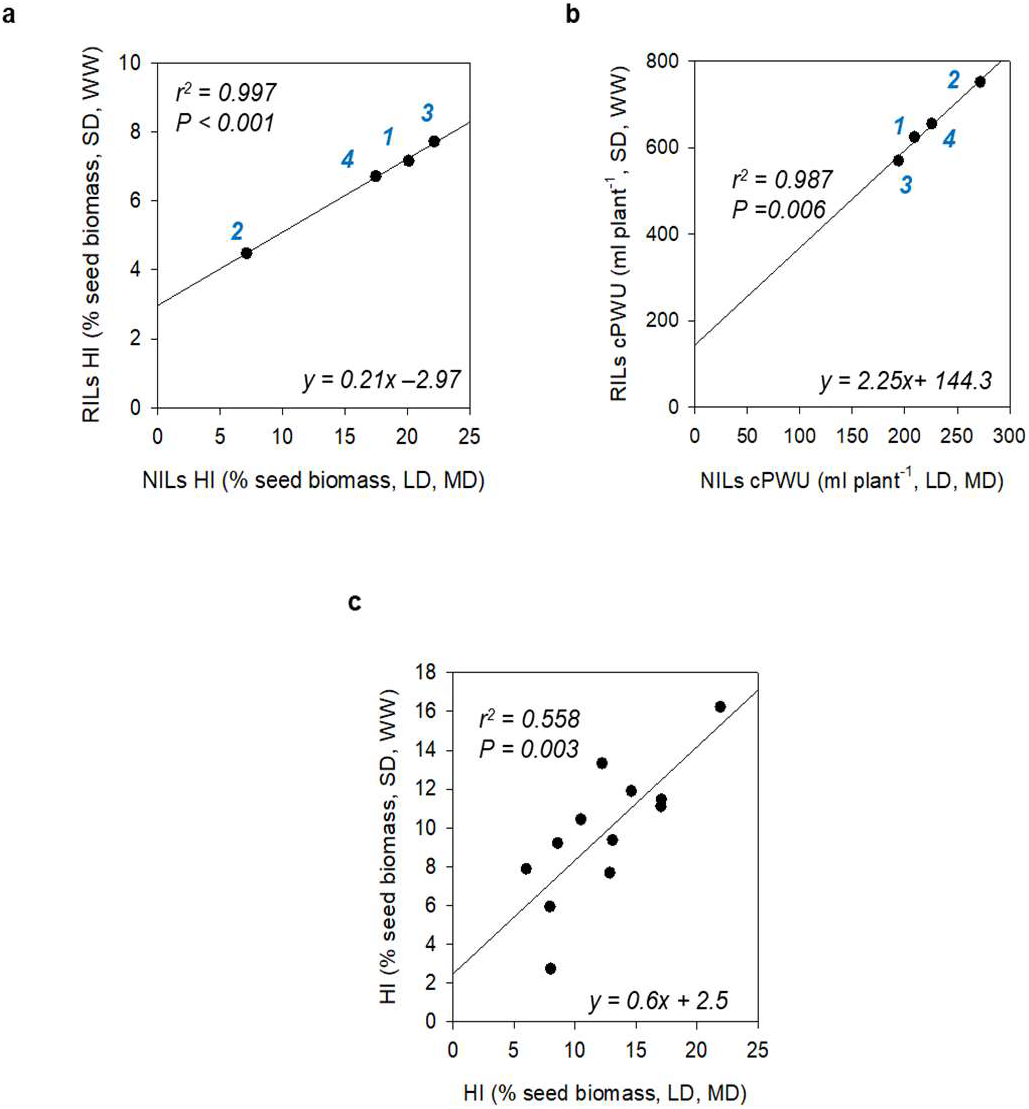
Comparison of water use parameters and HI parameters across different growth and watering regimes. **a** - correlation of Harvest index (HI) of the four *FRI/FLC* allelic groups tested in RILs and NILs. RILs were subjected to the growth regime shown in Figure 1A (SD, WW), while NILs were subjected to growth regime shown in Figure 1B (LD, MD), **b** - correlation between cPWU and cPWU of the four *FRI/FLC* allelic groups tested in RILs and NILs grown under two different day length and watering regimes (SD, WW and LD, MD), and **c** - correlation of Harvest index (HI) of 12 ecotypes subjected to the growth different growth regimes shown in Figure 1. The lines represent the equation of the linear regression model. The P-value of the slope parameter and adjusted R^2^ value associated with the linear model are provided for each association. SD-short day, LD – long day, WW – well watered, MD – moderate drought. Allelic combinations: 1 - *fri/FLC*, 2 - *FRI/FLC*, 3 – *fri/flc* and 4 – *FRI/flc*.

### Gene expression

The detected QTL regions contained many genes, as such we explored gene expression differences between the two parents within the mapping intervals for all three mapped traits. This was achieved using a publicly available microarray experiment comparing C24 and Col-0 (Bechtold *et al*. 2010) and RNAseq data of both parental accessions (Brosché *et al*. 2014). In total 9906 protein coding genes were identified within the 95% Bayesian credible intervals on chromosomes 4 and 5 (Table 3), of which 304 showed differential expressions between Col-0 and C24 (Tables S8, S9). We randomly selected three to four differentially expressed genes (up and down) for each interval, whilst also including *FRI*, *FLC* and *FLOWERING LOCUS T* (*FT*; Chromosome One) for analysis of gene expression in the NILs and both parental lines (Table S10) at 26- and 48-days post germination.

Early studies have shown that *FRI* up-regulates *FLC* expression in ecotypes that have the active allele of *FRI* (Michaels & Amasino 1999; Sheldon *et al*. 1999). NILs carrying the C24 *FRI* allele (Table S5) showed elevated *FLC* expression at 26- and 43-days post germination in plants grown under short-day controlled environment conditions (Fig. 1, 8a). Variation in *FLC* and *FRI* expression at 43 days post germination showed a significant association with flowering time and mPWU (Table S11), which was independent of *FT* expression (Table S11). This is in line with QTL mapping results where a significant association of the allelic state of *FRI* and *FLC* with flowering time and PWU was observed under short-day controlled environment conditions (Fig. 3a,b; Fig. 4; Fig. S8a,b). Other highly differentially expressed genes in the mapping intervals on chromosome 4 and 5 showed no specific pattern that significantly correlated with the flowering time phenotype or mPWU observed in the NILs across the two developmental stages (Table S11).

**Figure 8.**
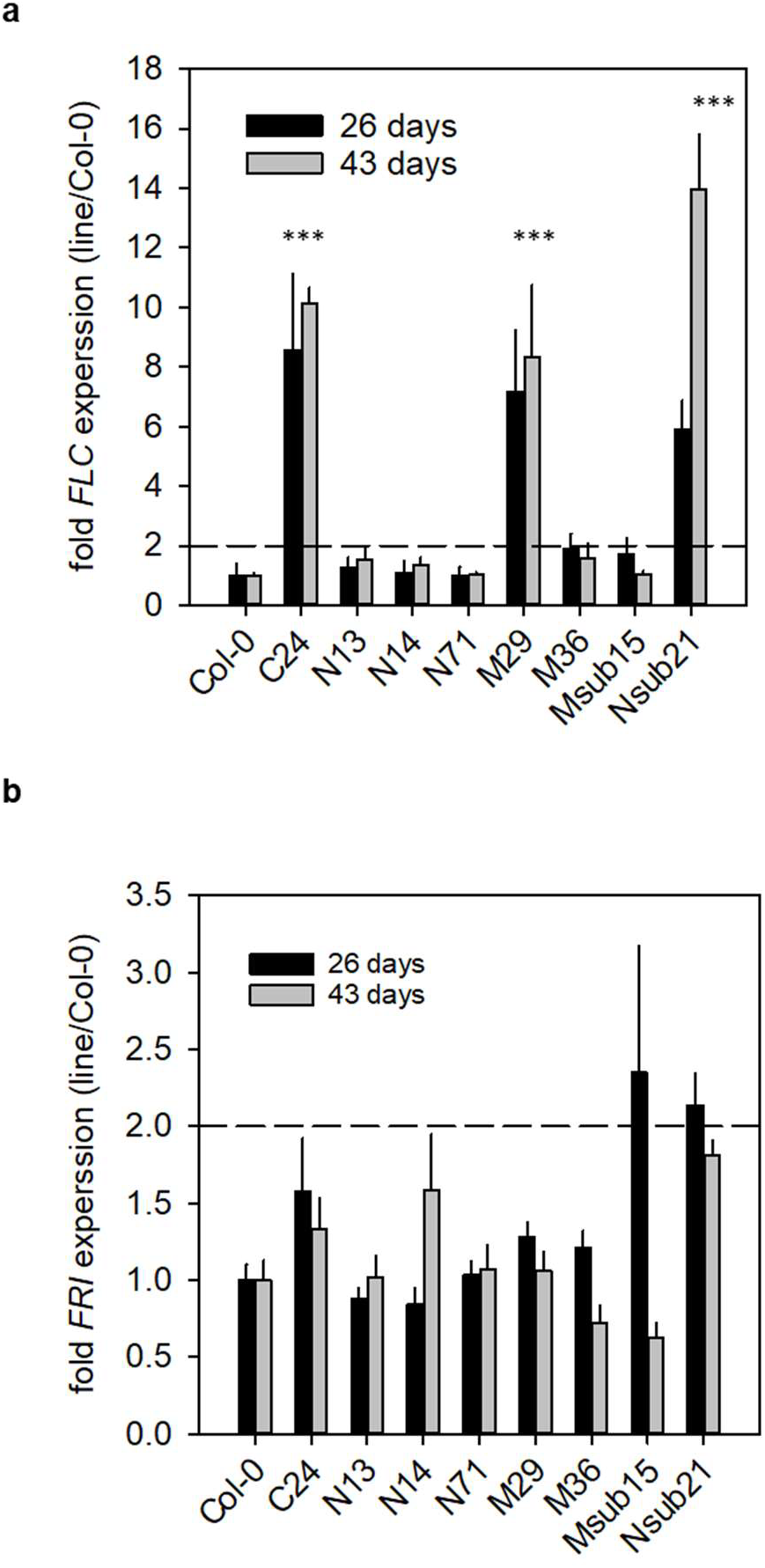
Expression of candidate genes in mapping interval. **a** – gene expression of *FLC* at 26 days after sowing (26 days) and 43 days after sowing (43 days). The stars above the columns denote significant different (P < 0.01) expression level compared to Col-0 at both time points. **b** - gene expression of *FRI* at 26 days after sowing (26 days) and 43 days after sowing (43 days). No significant gene expression levels compared between either the NILs or C24 and Col-0 were detected.

## Discussion

The ecotype C24 has an unusually rare combination of traits resulting in increased drought resistance, reduced VWU and increased WP (Bechtold *et al*. 2010; Ferguson *et al*. 2018), as well as resistance to a number of other abiotic and biotic stresses (Brosché *et al*. 2010; Lapin *et al*. 2012; Xu *et al*. 2015; Bechtold *et al*. 2018).

*WUE_i_* is considered to play a key role in plant water use (Steduto *et al*. 2007) as it relates equally to water loss by transpiration and net carbon gain, thus impacting on biomass production (Steduto *et al*., 2007; Long *et al*., 2015). Because of the relationship between leaf and plant-level *WUE* parameters, high leaf-level *WUE* is seen as an important trait for minimising water loss in many different plants species (Blum 2009; Sinclair & Rufty 2012; Vadez, Kholova, Medina, Kakkera & Anderberg 2014). In addition, *WUE* is often referred to as a drought adaptation trait (Condon *et al*. 2004; Comstock *et al*. 2005; McKay *et al*. 2008) because of the *A*/*g_s_* correlation, where *WUE* can increase during drought stress when stomata close, especially when A is not yet proportionally affected (Meinzer, Goldstein & Jaimes 1984; Gilbert, Holbrook, Zwieniecki, Sadok & Sinclair 2011; Easlon *et al*. 2014). However, WUE only evaluates how much water a plant needs to fix carbon, and in Arabidopsis, where within species variation in WUE is predominantly driven by variation in stomatal conductance (Easlon *et al*. 2014; Ferguson *et al*. 2018), overall plant water use will therefore be the main driver of *TE*.

### The importance of flowering time for plant water-use strategies

In natural populations, such as Arabidopsis, few studies have compared leaf-level measurements with whole-plant estimates of *WUE* (i.e. TE or *WP*; Bechtold *et al*. 2010, 2013; Easlon *et al*. 2014), and often leaf-level *WUE* measurements have been exploited as a screening tool to identify genes that could optimise water requirements and yield (McKay, Richards & Mitchell-Olds 2003; McKay *et al*. 2008; Hausmann *et al*. 2005; Juenger, Mckay, Hausmann, Keurentjes & Sen 2005; Masle *et al*. 2005). Natural genetic variation for δ^13^C has been demonstrated in Arabidopsis (Bouchabke-Coussa *et al*. 2008; Easlon *et al*. 2014; Kenney *et al*. 2014; Verslues & Juenger 2011), and QTL mapping has successfully elucidated the genetic basis of δ^13^C (Ghandilyan *et al*. 2009; Lovell *et al*. 2015; McKay *et al*. 2003; Hausmann *et al*. 2005; Juenger *et al*. 2005; Masle *et al*. 2005; McKay *et al*. 2008). Interestingly, a positive genetic correlation between flowering time and δ^13^C has been reported (McKay *et al*., 2003; Easlon *et al*., 2014), while other studies found a negative genetic correlation between flowering time and water content (Loudet *et al*., 2002, 2003). Despite these differences, the link between flowering time and plant water status is undeniable. Furthermore, natural polymorphisms of FRI and FLC have been identified as key determinants of the natural variation in δ^13^C (McKay *et al*. 2003, 2008; Kenney *et al*. 2014; Lovell *et al*. 2015), and FLC is also known to control the circadian rhythm of leaf movement (Edwards *et al*., 2006). It was therefore suggested that FLC may also regulate stomatal transpiration (Edwards *et al*., 2006), because accessions with a non-functional allele of FLC, showed reduced flowering time and increased water content (Loudet *et al*., 2002, 2003). Similarly, C24 possess a non-functional allele of FLC, exhibits a high relative water content (RWC) and low stomatal conductance (Bechtold *et al*., 2010; Fig. S10a,b). Our data suggests that although flowering time achieved through different combinations of weak or non-functional alleles of *FRI* and *FLC* explained most of the variation in plant water use (Fig. 4; Fig S8a,b), leaf-level traits associated with the lowered stomatal conductance phenotype were independent of variation at these genes (Fig S10). In addition, VWU, average daily water-use or the dehydration response were also not affected by the allelic combinations of *FRI* and *FLC* (Fig. S5c,d, Fig. 5d, Fig. S8c). Accordingly, QTLs identified for VWU did not overlap with the two major intervals containing *FRI* and *FLC* (Fig. 3c, Table 2). Importantly, plants with high mPWU also used more water on a daily basis, which suggests that life-time PWU is not only driven by flowering time but also by short-term water use strategies (Fig. 5c, Fig. S9).

In this study, cPWU and mPWU was clearly associated with increased flowering time (Fig. 2a). Mapping identified three QTLs for cPWU located on chromosomes 3, 4 and 5, and given the observed relationships between lifespan and water use (Fig. 2a), two also overlapped with flowering time QTLs (Fig. 3, Table 2). FRI and FLC were determined to be the causal genes underlying the overlapping QTLs on chromosomes 4 and 5 respectively (Fig. S1), which reinforced the role of flowering time in determining lifetime PWU. This is perhaps unsurprising, since a plant that lives for a longer period is likely to use more water, however this occurred without apparent gain of reproductive biomass (Fig. 6b; Fig. S12b). Interestingly, other development associated genes such as *ERECTA* (Masle *et al*., 2005; Villagarcia *et al*., 2012; Shen *et al*., 2015), *SHORT VEGETATIVE PROTEIN* (SVP or AGL22; Bechtold *et al*., 2016) and *HEAT SHOCK TRANSCRIPTION FACTOR A1b* (Bechtold *et al*., 2013; Albihlal *et al*., 2018), have been shown to affect stomatal function, stress tolerance and plant development in Arabidopsis and other plant species.

Similarly, the lack of a significant positive correlation between δ^13^C and flowering time in the NILs suggested that the variation in δ^13^C was independent of *FRI* and *FLC* in this mapping population (Fig S10c). However, increased δ^13^C coincided with reduced stomatal conductance and the distinctive growth phenotype of the C24 rosette (Fig. S10b,d). In Arabidopsis, δ^13^C is regulated by variation in stomatal conductance (Masle *et al*. 2005), which clearly corroborates the observed link between g_s_ and δ^13^C in the NILs and the independence from *FRI* and *FLC*. C24 is also more drought tolerant compared to Col-0 based on rosette wilting phenotypes after dehydration (Bechtold *et al*. 2010), and the drought response parameters were also independent of *FRI* and *FLC* in the RIL population (Fig. S11).

### The impact of day length on flowering time and water use

Col-0 is a rapid cycling ecotype (Shindo *et al*. 2005) and the higher *FLC* expression levels in C24 would suggest a late-flowering phenotype compared to Col-0 (Fig. 8a). However, early genetic studies have shown that C24 contains an allele of *FLC* that suppresses the late flowering phenotype caused by dominant alleles of *FRI*, whereas Col-0 contains an allele of *FLC* that does not suppress the late-flowering caused by dominant FRI alleles (Koornneef *et al*. 1994; Lee *et al*. 1994; Sanda & Amasino 1995). Therefore, we do not see a significant difference in flowering time between Col-0 and C24 in un-vernalised plants (Fig. 4b). The transition from short- to long-day conditions as part of our growing regimes (Fig. 1) mimics the natural progression in day length from spring to summer, which is commonly experienced by spring/summer annuals. Despite the difference in day length and watering regimes between the short dehydration and moderate drought treatments (Fig. 1), PWU and biomass allocation were significantly correlated between experiments (Fig. 7). This suggested that even though absolute values for HI and PWU were different the relative difference between lines remained the same (Fig. 7), indicating that day length does not alter overall water use and developmental strategies in a genotype-by-environment specific manner.

With respect to the above, it is worth noting that subjecting summer or winter annual ecotypes to long photoperiods may result in outcomes that could be problematic especially when assessing mechanisms related to leaf level *WUEi* drought resistance strategies, since these are often closely linked to flowering time. For example, Riboni *et al*. (2013, 2014) demonstrated that the induced drought escape mechanisms in Arabidopsis are promoted by the drought mediated up-regulation of florigens in an ABA- and photoperiod-dependent manner, so that early flowering (drought escape) can only occur under long days, independent of *FT* and *CONSTANS*. This is in line with our observation that flowering time and mPWU are associated with *FRI* and *FLC* expression, but seemingly independent of *FT* expression (Fig. 8., Tables S10, S11).

### The role of *FRI* and *FLC* in determining water use and biomass allocation

*FRI* and *FLC* respond to seasonal variation in temperature, thus play a crucial role in floral transitioning (Koornneef *et al*. 1994; Lee *et al*. 1994; Michaels & Amasino 2001). *FLC* is a MADS box transcription factor that inhibits the transition to flowering by repressing the expression of floral integrators, such as *FT* and *SUPPRESSOR OF OVEREXPRESSION OF CONSTANS1* (*SOC1*, Hepworth *et al*. 2002; Helliwell *et al*. 2006; Deng *et al*. 2011). Most rapid-cycling accessions of Arabidopsis contain naturally occurring loss-of-function mutations in FRI and therefore have low levels of *FLC* expression and are early flowering even in the absence of vernalization (Johanson *et al*. 2000).

Despite variation in cPWU mapping to *FLC* and *FRI*, we cannot explicitly rule out an indirect effect of flowering time differences on water use (Fig. S6c). Especially since *FLC* expression remained high in C24 and two NILs throughout the experiment (Fig. 8a), independent of the *FLC* allele present (Table S5). However, the reduction in mPWU attained via introgression of the non-functional Col-0 allele of *FLC* or the functional C24 *FRI* allele into the C24 and Col-0 genomic background respectively, demonstrates that although flowering time ultimately impacts PWU it does not confound the importance of these genes in determining PWU.

Interestingly, two major *FLC* haplogroups were associated with flowering time variation in *Arabidopsis* under field-like conditions, but only in the presence of functional *FRI* alleles (Caicedo, Stinchcombe, Olsen, Schmitt & Purugganan 2004). This is in line with our finding that the functional C24 allele of *FRI* (*FRI*) was required for increased *FLC* expression, even though *FRI* expression was not significantly altered (Fig. 8b, Table S10, S11). Furthermore, a study of ~150 accessions showed that the role of *FLC* in regulating flowering time is less important under short day conditions (Lempe *et al*. 2005), which suggests that the impact of *FLC* on PWU in our experiments may have been influenced by the environmental growth conditions such as photoperiod and potentially watering status (Fig. 1).

However, since *FLC* also acts in conjunction with other MADS-box proteins to regulate various aspects of plant development through a large variety of target genes (Deng *et al*. 2011), and rapid-cycling accessions contain a number of other genes regulating *FLC* expression, collectively known as the autonomous floral-promotion pathway (Michaels & Amasino 1999; Sheldon *et al*. 1999), we cannot rule out that other genetic factors affecting flowering time may indirectly contribute to the variation in whole plant water use.

The analysis of such putative relationships was beyond the scope of this study. Yet, the considerable number of *FLC* targets and their involvement in different developmental pathways may reflect an important strategy to integrate environmental signals and plant development to ensure reproductive success under many different conditions.

Short-term stress-mediated initiation of flowering pathways also involves the repression of *FLC* expression. Cold or saline stress-dependent activation of *miR169b*, was shown to repress the expression of the *NF-YA2* transcription factor, which in turn reduces *FLC* expression promoting early flowering (Xu *et al*. 2014). Here, stress treatments were shown to accelerate flowering (escape response) involving the above-described signalling cascade. We have previously demonstrated that the experimental watering regimes employed in this study (Figure 1), do not initiate a similar escape response in the progenitors of the mapping population and a number of other rapid cycling ecotypes (Ferguson *et al*. 2018; Bechtold *et al*. 2010; 2013). Heat sensitivity has been associated with late flowering haplotypes in vernalised plants, and *FLC* haplotypes resulting in late flowering showed reduced silique length, suggesting a negative correlation between flowering time and seed productivity (Bac-Molenaar *et al*. 2015). This negative correlation corroborates our findings, where late flowering RILs and NILs, produced less seed biomass and vice versa independent of photoperiod and watering conditions (Fig. 6b, S13b).

However, well-known work from the previous decade has demonstrated a pleiotropic link between flowering time and δ^13^C (WUE; McKay *et al*. 2003, Juenger *et al*. 2005). Similarly, phenotypic positive associations between flowering time and δ^13^C have been reported (Easlon *et al*. 2014, Kenney *et al*. 2014). It has therefore been suggested that functional alleles of *FRI* and *FLC* indirectly increase δ^13^C, suggesting that late flowering genotypes have greater WUE (McKay *et al*. 2003). The other referenced studies here support this notion in terms of flowering time and *WUE*, but not with respect to the allelic state of FRI and *FLC*. The identification that non-functional and weak alleles of *FRI* and *FLC* facilitate reduced water use and improved whole plant water use efficiency (Fig. 4, 6) challenges this previous work and illuminates the necessity to assess WUE at the whole plant and life-time level.

### The relationship between leaf-level and whole-plant measures of water use

Leaf level measures of *WUE*, taken during vegetative growth are not representative of whole plant measures such as *TE* or *WP* (Fig. S2a, b). This suggests that plants with improved δ^13^C and/or *WUE_i_* are not necessarily diverting additionally acquired photosynthates toward reproductive growth. In addition, our estimation of *TE* is clearly biased towards the final above ground biomass, neglecting root architecture. It is well established that both root depth and density play a major role in optimising water uptake depending on the hydrological conditions (Falik, Reides, Gersani & Novoplansky 2005; Czyz & Dexter 2012), but variation here may have been limited due to their likely pot bound nature. However, the relative performance of NILs and ecotypes was highly correlated between different experiments (Fig 7), suggesting that the variation observed for *TE* even though biased may reflect actual genotypic differences.

Different drought resistance mechanisms, such as avoidance by maintaining high plant water status and/or drought escape through early flowering (Levitt 1985) are critical from an ecological standpoint, facilitating population persistence in regions characterised by frequent and/or extended periods of reduced water availability (Araus, Slafer, Reynolds & Royo 2002; Gechev, Dinakar, Benina, Toneva & Bartels 2012; Kooyers 2015; Kooyers, Greenlee, Colicchio, Oh & Blackman 2015). However, leaf-level traits such as high *WUE_i_*/δ^13^C, aimed at preserving water may not always ensure high productivity, while lifespan also determines water use but not necessarily biomass production (Fig. 6; Fig. S12b), or allocation (Fig. S12a-c; Ferguson *et al*. 2018). In late flowering plants, photosynthates are not translocated to reproductive sinks, but instead to vegetative biomass (Figs. S2d), which either suggests poor resource allocation in late flowering ecotypes, or a diversion of resources toward abiotic stress defence mechanisms associated with reduced water availability (Claeys, Inze & Inzé 2013). Recent studies on the perennial species *Arabidopsis lyrata* and 35 Arabidopsis thaliana accessions highlighted that populations increased their reproductive output while reducing vegetative growth (Remington *et al*. 2013; Ferguson *et al*. 2018), which may be even more prevalent in annual plants that only have one opportunity at reproduction. While recent reports have clearly shown that there is a selection on early flowering in Arabidopsis due to increased plant fitness (Ågren, Oakley, Lundemo & Schemske 2017; Austen, Rowe, Stinchcombe & Forrest 2017; Gnan, Marsh & Kover 2017), still little is known about the genotype-to-phenotype basis of this resource allocation trade-off.

## Conclusion

We conclude that flowering is the predominant determinant of lifetime PWU strategies, a critical life history trait that is important for seed production. Absolute water use at the vegetative growth stage contributes to overall PWU, albeit to a much-reduced degree. The causal genes that underlie these QTLs are ambiguous and will require further fine-mapping. We have demonstrated that Arabidopsis plant water use strategies are independent of traditional leaf-level measures of drought tolerance, *WUE* and biomass traits, and consequently genes identified based on these traditional performance traits may not lead to improved productivity under water limiting or water-replete conditions.

## Acknowledgements

We thank Susan Corbett and Philip M Mullineaux for help with the continuous moderate drought experiments. J.N.F was funded by a BBSRC CASE award (BB/J012564/1). U.B. is supported by the University of Essex. J.N.F. and U.B. performed and analysed all experiments. O.B and R.M provided and analysed plant material, including the RIL and NIL populations. U.B., J.N.F., and M.H. planned and designed the experiments. U.B. and J.N.F wrote the manuscript with the input from all authors. Isotopic measurements were performed by C. Hossann at the Plateforme Technique d’Ecologie Fonctionnelle (PTEF) (OC 081, INRA Nancy, France).

## Supporting Information

**Figure S1. The single nucleotide polymorphism (SNP) markers used and their position on the re-estimated linkage map. a** - *InDel* markers for *FRI* and *FLC*, used to score the C24 x Col-0 RIL population, and **b** - Position in cMs of all markers on the re-estimated genetic map.

**Figure S2. Comparison of leaf level water use efficiency and biomass level water use efficiency parameters. a - b** Relationship between δ^13^C, and whole plant water use efficiency parameters biomass level *WUE* parameters: *TE* (transpiration efficiency) and *WP* (water productivity) and **c – d** Relationship between *WUE_i_*, and whole plant water use efficiency parameters biomass level *WUE* parameters, *TE* and *WP*. The associations are not significant in all cases.

**Figure S3. Distribution of estimated means for all traits assessed as part of the QTL mapping. a** - vegetative water use (VWU), **b** - days to flowering, **c** - seed biomass, **d** - calculated lifetime plant water-use (cPWU), **e** - dehydration plasticity (VWU plasticity), and **f** - breakpoint (rSWC) of the segmented regression. For all traits, a Shaprio-Wilk test of normality was performed on the estimated means of all RILs, where all traits demonstrated variation that was not significantly different from a normal distribution (P > 0.05). Green arrows indicate the position of C24 and red arrows indicate the position of Col-0. The estimated means for the parental lines are also provided (Red – Col-0, Green – C24)

**Figure S4. Comparison of different methods for mapping QTL for cPWU. Black** – Hayley-Knott method. **Red** – Expectation-Maximization method. **Blue** – Multiple-Imputation method.

**Figure S5: Further QTL mapping results. a** - LOD profiles for seed biomass, with no significant QTL detected, **b** - LOD profiles for dehydration plasticity, with no significant QTL detected, **c** - LOD profiles for breakpoint (rSWC), with no significant QTL detected, and **d** – LOD profiles for slope 1, with one significant QTL detected. The dashed horizontal red line indicates the 0.05 genome-wide significance threshold.

**Figure S6: Single QTL mapping for calculated plant water use with and without traits as covariates. a** – Without a trait covariate. **b** – With rosette biomass as a trait covariate. **c** – With flowering time as a trait covariate. **d**-With vegetative water use as a covariate.

**Figure S7: LOD scores for a two dimensional genome scan for calculated plant water use.** Values in the upper left triangle represent the full QTL model. Values on the lower right triangle represent the likelihood ratio comparing the full model with QTLs on all chromosomes with the single QTL model, thus indicating the presence of epistatic interactions.

**Figure S8: Trait performances of genotypes harbouring different allelic combinations of the FRIGIDA (FRI) and FLOWERING LOCUS C (FLC) genes.** Boxplots describing the variation for traits assessed for the 4 groups based on allelic combination of FRI and FLC, **a** – mPWU in the NILs, **b** - days to flowering in the NILs, **c** - VWU based on allelic combinations of FRI/FLC in the RILs, and **d** - VWU based on allelic combinations of FRI/FLC in the NILs. The letters (a, b, and c) above the boxplot denote the post-hoc Tukey groups, where allelic groups whose letters are different are significantly different from one another for that particular trait at P < 0.05. The bold line in the centre of the boxplots represents the median, the box edges represent the 25th (lower) and 75th (upper) percentiles, the whiskers extend to the most extreme data points that are no more than 1.5x the length of the upper or lower segment. Outliers are data points that lie outside the 1.5x interquartile range both above the upper quartile and below the lower quartile.

**Figure S9: The contribution of mean daily water use in the 12 ecotypes. a** - relationship between flowering time and mean daily water use, **b** - relationship between rosette biomass and mean daily water use, and **c** - relationship between mean daily water use and mPWU. The linear model of the relationship between mean long-term water use and mean daily water use is provided. R2 and P values are provided where a significant relationship was identified.

**Figure S10: Phenotype of NILs and parental lines. a** - boxplots of leaf level WUE (δ^13^C) for the 4 groups based on allelic combination of both *FRI* and *FLC* in the NILs and both parents. The letters (a, b) denote the post-hoc Games-Howell groups, where allelic groups whose letters are different are significantly different from one another for that trait at P < 0.05. The bold line in the centre of the boxplots represents the median, the box edges represent the 25th (lower) and 75th (upper) percentiles, the whiskers extend to the most extreme data points that are no more than 1.5x the length of the upper or lower segment. Outliers are data points that lie outside the 1.5x interquartile range both above the upper quartile and below the lower quartile, b - phenotype scoring based on rosette growth (panel C), stomatal conductance (*gs*) and δ^13^C measurements. There was a significant negative correlation between gs and δ^13^C. r^2^= 0.781, P < 0.001, **c** - relationship between δ^13^C and flowering time, and **d** - rosette growth at 25 days post sowing.

**Figure S11: Boxplots of drought response parameters derived from segmented regression analysis based on allelic combinations of FRI/FLC. a** - dehydration plasticity (see Table 1), and **b** - breakpoint (rSWC) between segment 1 and 2. Both parameters were calculated using predicted means of the short dehydration experiment performed on the RIL population. No significant differences were detected between the four allelic combinations. The bold line in the centre of the boxplots represents the median, the box edges represent the 25^th^ (lower) and 75^th^ (upper) percentiles, the whiskers extend to the most extreme data points that are no more than 1.5x the length of the upper or lower segment. Outliers are data points that lie outside the 1.5x interquartile range both above the upper quartile and below the lower quartile.

**Figure S12: Boxplots of biomass parameters based on allelic combinations of FRI/FLC in the NILs a** – above ground biomass, **b** – seed biomass, and **c** – rosette biomass. The letters (a, b, and c) above the boxplot denote the post-hoc Tukey groups, where allelic groups whose letters are different are significantly different from one another for that trait at P < 0.05. The bold line in the centre of the boxplots represents the median, the box edges represent the 25th (lower) and 75th (upper) percentiles, the whiskers extend to the most extreme data points that are no more than 1.5x the length of the upper or lower segment. Outliers are data points that lie outside the 1.5x interquartile range both above the upper quartile and below the lower quartile.

**Figure S13: Above ground biomass allocation. a** - biomass distribution in the NILs of moderate drought stressed plants. **b** - biomass distribution in 164 RILs including both parents.

**Table S1:** Ecotypes used in benchmarking experiment

**Table S2:** RIL genotypes according to Tjörék *et al*. (2006)

**Table S3:** Primers used in genotyping and qPCR

**Table S4:** Genotyping of FRI and FLC alleles in RIL population using InDel markers, scored by qPCR and high-resolution melt (HRM) curve.

**Table S5:** Genotypes of near isogenic lines (NILs)

**Table S6:** Correlation matrix of traits analysed for the 12 ecotypes population

**Table S7:** Correlation matrix of traits analysed for the RIL population

**Table S8:** Number of differentially expressed protein coding genes in mapping intervals

**Table S9:** IDs of differentially expressed genes in mapping intervals

**Table S10:** Fold expression and error (Line/Col-0) of selected DE genes in three mapping intervals at 26- and 43 days post germination (n= 3).

**Table S11:** Association between gene expression and mPWU and flowering time (Flowering). Genes FLOWERING LOCUS T (FT), FRI, FLC and At4g00960.

## References

Ågren J., Oakley C.G., Lundemo S. & Schemske D.W. (2017) Adaptive divergence in flowering time among natural populations of Arabidopsis thaliana: Estimates of selection and QTL mapping. Evolution 71, 550-564.

Van Aken O., De Clercq I., Ivanova A., Law S.R., Van Breusegem F., Millar A.H. & Whelan J. (2016) Mitochondrial and Chloroplast Stress Responses Are Modulated in Distinct Touch and Chemical Inhibition Phases. Plant physiology 171, 2150-65.

Anderson J.T. (2016) Plant fitness in a rapidly changing world. New Phytologist 210, 81-87.

Araus J.L., Slafer G. a., Reynolds M.P. & Royo C. (2002) Plant breeding and drought in C3 cereals: What should we breed for? Annals of Botany 89, 925-940.

Austen E.J., Rowe L., Stinchcombe J.R. & Forrest J.R.K. (2017) Explaining the apparent paradox of persistent selection for early flowering. New Phytologist 215, 929-934.

Bac-Molenaar J.A., Fradin E.F., Becker F.F.M., Rienstra J.A., van der Schoot J., Vreugdenhil D. & Keurentjes J.J.B. (2015) Genome-Wide Association Mapping of Fertility Reduction upon Heat Stress Reveals Developmental Stage-Specific QTLs in Arabidopsis thaliana. The Plant Cell 27, 1857-1874.

Bechtold U., Albihlal W.S., Lawson T., Fryer M.J., Sparrow P.A.C., Richard F.F., … Mullineaux P.M. (2013) Arabidopsis HEAT SHOCK TRANSCRIPTION FACTORA1b overexpression enhances water productivity, resistance to drought, and infection. Journal of Experimental Botany 64, 3467-3481.

Bechtold U., Lawson T., Mejia-Carranza J., Meyer R.C., Brown I.R., Altmann T., … Mullineaux P.M. (2010) Constitutive salicylic acid defences do not compromise seed yield, drought tolerance and water productivity in the Arabidopsis accession C24. Plant, Cell and Environment 33, 1959-1973.

Bechtold U., Penfold C.A., Jenkins D.J., Legaie R., Moore J.D., Lawson T., … Mullineaux P.M. (2016) Time-Series Transcriptomics Reveals That AGAMOUS-LIKE22 Affects Primary Metabolism and Developmental Processes in Drought-Stressed Arabidopsis. The Plant Cell 28, 345-66.

Bechtold U., Richard O., Zamboni A., Gapper C., Geisler M., Pogson B., … Mullineaux P.M. (2008) Impact of chloroplastic- and extracellular-sourced ROS on high light-responsive gene expression in Arabidopsis. Journal of Experimental Botany 59, 121-133.

Blum A. (2005) Drought resistance, water-use efficiency, and yield potential—are they compatible, dissonant, or mutually exclusive? Australian Journal of Agricultural Research 56, 1159-1168.

Blum A. (2009) Effective use of water (EUW) and not water-use efficiency (WUE) is the target of crop yield improvement under drought stress. Field Crops Research 112, 119-123.

Bouchabke-Coussa O., Quashie M.-L., Seoane-Redondo J., Fortabat M.-N., Gery C., Yu A., … Durand-Tardif M. (2008) ESKIMO1 is a key gene involved in water economy as well as cold acclimation and salt tolerance. BMC plant biology 8, 125.

Boyes D.C., Zayed A.M., Ascenzi R., Mccaskill A.J., Hoffman N.E., Davis K.R. & Görlach J. (2001) Growth Stage - Based Phenotypic Analysis of Arabidopsis: A Model for High Throughput Functional Genomics in Plants. 13, 1499-1510.

Brendel O., Le Thiec D., Scotti-Saintagne C., Bod◌้n◌่s C., Kremer A. & Guehl J.M. (2008) Quantitative trait loci controlling water use efficiency and related traits in Quercus robur L. Trees Genetics & Genomes.

Broman K.W. (2009) A brief tour of R / qtl. 889-890.

Broman K.W., Wu H., Sen S. & Churchill G.A. (2003) R/qtl: QTL mapping in experimental crosses. Bioinformatics 19, 889-890.

Brosché M., Merilo E., Mayer F., Pechter P., Puzõrjova I., Brader G., … Kollist H. (2010) Natural variation in ozone sensitivity among Arabidopsis thaliana accessions and its relation to stomatal conductance. Plant, Cell and Environment 33, 914-925.

Caicedo A.L., Stinchcombe J.R., Olsen K.M., Schmitt J. & Purugganan M.D. (2004) Epistatic interaction between Arabidopsis FRI and FLC flowering time genes generates a latitudinal cline in a life history trait. Proceedings of the National Academy of Sciences.

Cernusak L. a, Winter K. & Turner B.L. (2009) Physiological and isotopic (delta(13)C and delta(18)O) responses of three tropical tree species to water and nutrient availability. Plant, cell & environment 32, 1441-55.

Cernusak L.A., Ubierna N., Winter K., Holtum J.A.M., Marshall J.D. & Farquhar G.D. (2013) Environmental and physiological determinants of carbon isotope discrimination in terrestrial plants. New Phytologist 200, 950-965.

Claeys H., Inze D. & Inzé D. (2013) The Agony of Choice: How Plants Balance Growth and Survival under Water-Limiting Conditions. PLANT PHYSIOLOGY 162, 1768-1779.

Comstock J.P., McCouch S.R., Martin B.C., Tauer C.G., Vision T.J., Xu Y. & Pausch R.C. (2005) The effects of resource availability and environmental conditions on genetic rankings for carbon isotope discrimination during growth in tomato and rice. Functional Plant Biology 32, 1089-1105.

Condon A.G., Richards R.A., Rebetzke G.J. & Farquhar G.D. (2004) Breeding for high water-use efficiency. Journal of Experimental Botany 55, 2447-2460.

Craig H. (1957) Isotopic standards for carbon and oxygen and correction factors for mass-spectrometric analysis of carbon dioxide. Geochimica et Cosmochimica Acta 12, 133-

Czyz E.A. & Dexter A.R. (2012) Plant wilting can be caused either by the plant or by the soil. Soil Research 50, 708-713.

Deng W., Ying H., Helliwell C.A., Taylor J.M., Peacock W.J. & Dennis E.S. (2011) FLOWERING LOCUS C (FLC) regulates development pathways throughout the life cycle of Arabidopsis. Proceedings of the National Academy of Sciences.

Donovan L.A., Dudley S.A., Rosenthal D.M. & Ludwig F. (2007) Phenotypic selection on leaf water use efficiency and related ecophysiological traits for natural populations of desert sunflowers. Oecologia.

Easlon H.M., Nemali K.S., Richards J.H., Hanson D.T., Juenger T.E. & McKay J.K. (2014) The physiological basis for genetic variation in water use efficiency and carbon isotope composition in Arabidopsis thaliana. Photosynthesis Research 119, 119-129.

Von Euler T., Ågren J. & Ehrlén J. (2014) Environmental context influences both the intensity of seed predation and plant demographic sensitivity to attack. Ecology 95, 495-504.

Falik O., Reides P., Gersani M. & Novoplansky A. (2005) Root navigation by self inhibition. Plant, Cell and Environment 28, 562-569.

Farquhar G.D. & Von Caemmerer S. (1982) Modelling of photosynthetic response to environmental conditions. In Encyclopedia of plant physiology. pp. 549-587.

Farquhar G.D., Ehleringer J.R. & Hubick K.T. (1989) Carbon Isotope Discrimination and Photosynthesis. Annual Review of Plant Physiology and Plant Molecular Biology 40, 503-537.

Ferguson J.N., Humphry M., Lawson T., Brendel O. & Bechtold U. (2018) Natural variation of life-history traits, water use, and drought responses in Arabidopsis. Plant Direct 2, e00035.

Gazzani S., Gendall A.R., Lister C. & Dean C. (2003) Analysis of the Molecular Basis of Flowering Time Variation in Arabidopsis Accessions. Plant Physiology 132, 1107-1114.

Gechev T.S., Dinakar C., Benina M., Toneva V. & Bartels D. (2012) Molecular mechanisms of desiccation tolerance in resurrection plants. Cellular and Molecular Life Sciences 69, 3175-3186.

Ghandilyan A., Ilk N., Hanhart C., Mbengue M., Barboza L., Schat H., … Aarts M.G.M. (2009) A strong effect of growth medium and organ type on the identification of QTLs for phytate and mineral concentrations in three arabidopsis thaliana RIL populations. Journal of Experimental Botany 60, 1409-1425.

Gilbert M.E., Holbrook N.M., Zwieniecki M.A., Sadok W. & Sinclair T.R. (2011) Field confirmation of genetic variation in soybean transpiration response to vapor pressure deficit and photosynthetic compensation. Field Crops Research 124, 85-92.

Gnan S., Marsh T. & Kover P.X. (2017) Inflorescence photosynthetic contribution to fitness releases Arabidopsis thaliana plants from trade-off constraints on early flowering. PLoS ONE 12.

Granier C. & Tardieu F. (2009) Multi-scale phenotyping of leaf expansion in response to environmental changes: The whole is more than the sum of parts. Plant, Cell and Environment 32, 1175-1184.

Gruber F., Falkner F.G., Dorner F. & Hämmerle T. (2001) Quantitation of Viral DNA by Real-Time PCR Applying Duplex Amplification, Internal Standardization, and Two-Color Fluorescence Detection. Applied and Environmental Microbiology 67, 2837-2839.

Haley C.S. & Knott S.A. (1992) A simple regression method for mapping quantitative trait loci in line crosses using flanking markers. Heredity 69, 315-24.

Halperin O., Gebremedhin A., Wallach R. & Moshelion M. (2017) High-throughput physiological phenotyping and screening system for the characterization of plant???environment interactions. Plant Journal 89, 839-850.

Hausmann N.J., Juenger T.E., Sen S., Stowe K. a., Dawson T.E., Simms E.L., … Simms E.L. (2005) Quantitative trait loci affecting δ13C and response to differential water availibility in Arabidopsis thaliana. Evolution 59, 81-96.

Helliwell C.A., Wood C.C., Robertson M., James Peacock W. & Dennis E.S. (2006) The Arabidopsis FLC protein interacts directly in vivo with SOC1 and FT chromatin and is part of a high-molecular-weight protein complex. Plant Journal 46, 183-192.

Hepworth S.R., Valverde F., Ravenscroft D., Mouradov A. & Coupland G. (2002) Antagonistic regulation of flowering-time gene SOC1 by CONSTANS and FLC via separate promoter motifs. EMBO Journal.

Johanson U., West J., Lister C., Michaels S., Amasino R. & Dean C. (2000) Molecular Analysis of FRÍGIDA, a Major Determinant of Natural Variation in Arabidopsis Flowering Time. Science 290, 344-348.

Juenger T.E., Mckay J.K., Hausmann N., Keurentjes J.J.B. & Sen S. (2005) Identification and characterization of QTL underlying whole-plant physiology in Arabidopsis thaliana: d 13 C, stomatal conductance and transpiration efficiency. Plant Cell and Environment 28, 697-708.

Kenney A.M., Mckay J.K., Richards J.H. & Juenger T.E. (2014) Direct and indirect selection on flowering time, water-use efficiency (WUE, ??13C), and WUE plasticity to drought in Arabidopsis thaliana. Ecology and Evolution 4, 4505-4521.

Koornneef M., Blankestijn???de Vries H., Hanhart C., Soppe W. & Peeters T. (1994) The phenotype of some late???flowering mutants is enhanced by a locus on chromosome 5 that is not effective in the Landsberg erecta wild???type. The Plant Journal 6, 911-919.

Kooyers N.J. (2015) The evolution of drought escape and avoidance in natural herbaceous populations. Plant Science 234, 155-162.

Kooyers N.J., Greenlee A.B., Colicchio J.M., Oh M. & Blackman B.K. (2015) Replicate altitudinal clines reveal that evolutionary flexibility underlies adaptation to drought stress in annual Mimulus guttatus. New Phytologist 206, 152-165.

Lander E.S. & Green P. (1991) Counting algorithms for linkage: correction to Morton and Collins. Annals of Human Genetics 55, 33-38.

Lapin D., Meyer R.C., Takahashi H., Bechtold U. & Van den Ackerveken G. (2012) Broad-spectrum resistance of Arabidopsis C24 to downy mildew is mediated by different combinations of isolate-specific loci. New Phytologist 196, 1171-1181.

Lee I., Michaels S.D., Masshardt A.S. & Amasino R.M. (1994) The late???flowering phenotype of FRIGIDA and mutations in LUMINIDEPENDENS is suppressed in the Landsberg erecta strain of Arabidopsis. The Plant Journal 6, 903-909.

Lempe J., Balasubramanian S., Sureshkumar S., Singh A., Schmid M. & Weigel D. (2005) Diversity of flowering responses in wild Arabidopsis thaliana strains. PLoS Genetics 1, 0109-0118.

Levitt J. (1985) Relationship of dehydration rate to drought avoidance, dehydration tolerance and dehydration avoidance of cabbage leaves, and to their acclimation during drought-induced water stress. Plant, Cell & Environment 8, 287-296.

Long S.P., Marshall-Colon A. & Zhu X.G. (2015) Meeting the global food demand of the future by engineering crop photosynthesis and yield potential. Cell 161, 56-66.

Lovell J.T., Mullen J.L., Lowry D.B., Awole K., Richards J.H., Sen S., … McKay J.K. (2015) Exploiting Differential Gene Expression and Epistasis to Discover Candidate Genes for Drought-Associated QTLs in Arabidopsis thaliana. The Plant Cell 27, 969-983.

Lynch M. & Walsh B. (1998) Genetics and analysis of quantitative traits.

Manichaikul A., Moon J.Y., Sen S., Yandell B.S. & Broman K.W. (2009) A model selection approach for the identification of quantitative trait loci in experimental crosses, allowing epistasis. Genetics 181, 1077-86.

Marguerit E., Bouffier L., Chancerel E., Costa P., Lagane F., -Marc Guehl J., … Brendel O. (2014) The genetics of water-use efficiency and its relation to growth in maritime pine. Journal of Experimental Botany 65, 4757-4768.

Masle J., Gilmore S.R. & Farquhar G.D. (2005) The ERECTA gene regulates plant transpiration efficiency in Arabidopsis. Nature 436, 866-870.

McKay J.K., Richards J.H. & Mitchell-Olds T. (2003) Genetics of drought adaptation in Arabidopsis thaliana: I. Pleiotropy contributes to genetic correlations among ecological traits. Molecular Ecology 12, 1137-1151.

McKay J.K., Richards J.H., Nemali K.S., Sen S., Mitchell-Olds T., Boles S., … Juenger T.E. (2008) Genetics of drought adaptation in Arabidopsis thaliana II. QTL analysis of a new mapping population, Kas-1 ?? Tsu-1. Evolution 62, 3014-3026.

Medrano H., Tom?s M., Martorell S., Flexas J., Hern?ndez E., Rossell? J., … Bota J. (2015) From leaf to whole-plant water use efficiency (WUE) in complex canopies: Limitations of leaf WUE as a selection target. Crop Journal 3, 220-228.

Meinzer F., Goldstein G. & Jaimes M. (1984) The effect of atmospheric humidity on stomatal control of gas exchange in two tropical coniferous species. Canadian Journal of Botany 62, 591-595.

Michaels S.D. (1999) FLOWERING LOCUS C Encodes a Novel MADS Domain Protein That Acts as a Repressor of Flowering. THE PLANT CELL ONLINE 11, 949-956.

Michaels S.D. & Amasino R.M. (1999) FLOWERING LOCUS C encodes a novel MADS domain protein that acts as a repressor of flowering. The Plant cell 11, 949-56.

Michaels S.D. & Amasino R.M. (2001) Loss of FLOWERING LOCUS C activity eliminates the late-flowering phenotype of FRIGIDA and autonomous pathway mutations but not responsiveness to vernalization. The Plant cell 13, 935-41.

Michaels S.D., He Y., Scortecci K.C. & Amasino R.M. (2003) Attenuation of FLOWERING LOCUS C activity as a mechanism for the evolution of summer-annual flowering behavior in Arabidopsis. Proceedings of the National Academy of Sciences of the United States of America 100, 10102-7.

Monclus R., Dreyer E., Villar M., Delmotte F.M., Delay D., Petit J.M., … Brignolas F. (2006) Impact of drought on productivity and water use efficiency in 29 genotypes of Populus deltoides x Populus nigra. New Phytologist 169, 765-777.

Monneveux P., Sánchez C., Beck D. & Edmeades G.O. (2006) Drought tolerance improvement in tropical maize source populations: Evidence of progress. Crop Science 46, 180-191.

Morison J.I.L., Baker N.R., Mullineaux P.M. & Davies W.J. (2008) Improving water use in crop production. Philosophical Transactions of the Royal Society of London. Series B, Biological Sciences 363, 639-658.

Muggeo M.V.M.R. (2017) Package ‘ segmented.’ Biometrika 58, 525-534.

Oliveros J.C. (2007) VENNY. An interactive tool for comparing lists with Venn Diagrams. BioinfoGP of CNB-CSIC, http://bioinfogp.cnnb.csic.es/tools/venny/index.ht.

Parry M.A.J., Flexas J. & Medrano H. (2005) Prospects for crop production under drought: Research priorities and future directions. Annals of Applied Biology 147, 211-226.

Parsons R., Weyers J.D.B., Lawson T. & Godber I.M. (1997) Rapid and straightforward estimates of photosynthetic characteristics using a portable gas exchange system. Photosynthetica 34, 265-279.

Passioura J. (2007) The drought environment: Physical, biological and agricultural perspectives. Journal of Experimental Botany 58, 113-117.

R Core Team (2015) R Development Core Team. R: A Language and Environment for Statistical Computing 55, 275-286.

Remington D.L., Leinonen P.H., Leppälä J. & Savolainen O. (2013) Complex genetic effects on early vegetative development shape resource allocation differences between Arabidopsis lyrata populations. Genetics 195, 1087-1102.

Riboni M., Galbiati M., Tonelli C. & Conti L. (2013) GIGANTEA Enables Drought Escape Response via Abscisic Acid-Dependent Activation of the Florigens and SUPPRESSOR OF OVEREXPRESSION OF CONSTANS1. PLANT PHYSIOLOGY 162, 1706-1719.

Riboni M., Robustelli Test A., Galbiati M., Tonelli C. & Conti L. (2014) Environmental stress and flowering time: the photoperiodic connection. Plant signaling & behavior 9, e29036.

Roussel M., Dreyer E., Montpied P., Le-Provost G., Guehl J.M. & Brendel O. (2009) The diversity of 13C isotope discrimination in a Quercus robur full-sib family is associated with differences in intrinsic water use efficiency, transpiration efficiency, and stomatal conductance. Journal of Experimental Botany 60, 2419-2431.

Sanda S.L. & Amasino R.M. (1996) Ecotype-Specific Expression of a Flowering Mutant Phenotype in Arabidopsis thaliana. Plant physiology 111,641-644.

Seibt U., Rajabi A., Griffiths H. & Berry J.A. (2008) Carbon isotopes and water use efficiency: Sense and sensitivity. Oecologia 155, 441-454.

Sheldon C.C., Burn J.E., Perez P.P., Metzger J., Edwards J. a, Peacock W.J. & Dennis E.S. (1999) The FLF MADS Box Gene: A Repressor of Flowering in Arabidopsis Regulated by Vernalization and Methylation. THE PLANT CELL ONLINE 11,445-458.

Shindo C., Aranzana M.J., Lister C., Baxter C., Nicholls C., Nordborg M. & Dean C. (2005) Role of FRIGIDA and FLOWERING LOCUS C in Determining Variation in Flowering Time of Arabidopsis. Plant Physiology 138, 1163-1173.

Sinclair T.R. & Rufty T.W. (2012) Nitrogen and water resources commonly limit crop yield increases, not necessarily plant genetics. Global Food Security 1, 94-98.

Sletvold N. & Ågren J. (2015) Climate-dependent costs of reproduction: Survival and fecundity costs decline with length of the growing season and summer temperature. Ecology Letters 18, 357-364.

Törjék O., Witucka-Wall H., Meyer R.C., Von Korff M., Kusterer B., Rautengarten C., … Altmann T. (2006) Segregation distortion in Arabidopsis C24/Col-0 and Col-0/C24 recombinant inbred line populations is due to reduced fertility caused by epistatic interaction of two loci. Theoretical and Applied Genetics 113, 1551-1561.

Tisné S., Serrand Y., Bach L., Gilbault E., Ben Ameur R., Balasse H., … Loudet O. (2013) Phenoscope: An automated large-scale phenotyping platform offering high spatial homogeneity. Plant Journal 74, 534-544.

Törjék O., Meyer R.C., Zehnsdorf M., Teltow M., Strompen G., Witucka-Wall H., … Altmann T. (2008) Construction and analysis of 2 reciprocal arabidopsis introgression line populations. Journal of Heredity.

Vadez V., Kholova J., Medina S., Kakkera A. & Anderberg H. (2014) Transpiration efficiency: New insights into an old story. Journal of Experimental Botany 65, 6141-6153.

Verslues P.E. & Juenger T.E. (2011) Drought, metabolites, and Arabidopsis natural variation: A promising combination for understanding adaptation to water-limited environments. Current Opinion in Plant Biology 14, 240-245.

Xu E., Vaahtera L., Hõrak H., Hincha D.K., Heyer A.G. & Brosché M. (2015) Quantitative trait loci mapping and transcriptome analysis reveal candidate genes regulating the response to ozone in Arabidopsis thaliana. Plant, Cell and Environment 38, 1418-1433.

Xu M.Y., Zhang L., Li W.W., Hu X.L., Wang M.B., Fan Y.l Wang L. (2014) Stress-induced early flowering is mediated by miR169 in Arabidopsis thaliana. Journal of Experimental Botany.

Anderson J.T. (2016) Plant fitness in a rapidly changing world. New Phytologist 210, 81-87.

Broman K.W. (2009) A brief tour of R / qtl. 889-890.

Brosché M., Merilo E., Mayer F., Pechter P., Puzorjova I., Brader G., … Kollist H. (2010) Natural variation in ozone sensitivity among Arabidopsis thaliana accessions and its relation to stomatal conductance. Plant, Cell and Environment 33, 914-925.

Chen M. & Penfield S. (2018) Feedback regulation of COOLAIR expression controls seed dormancy and flowering time. Science.

Craig H. (1957) Isotopic standards for carbon and oxygen and correction factors for mass-spectrometric analysis of carbon dioxide. Geochimica et Cosmochimica Acta 12, 133-149.

Gruber F., Falkner F.G., Dorner F. & Hämmerle T. (2001) Quantitation of Viral DNA by RealTime PCR Applying Duplex Amplification, Internal Standardization, and Two-Color Fluorescence Detection. Applied and Environmental Microbiology 67, 2837-2839.

Lynch M. & Walsh B. (1998) Genetics and analysis of quantitative traits.

Masle J., Gilmore S.R. & Farquhar G.D. (2005) The ERECTA gene regulates plant transpiration efficiency in Arabidopsis. Nature 436, 866–870.

Muggeo M.V.M.R. (2017) Package ‘ segmented.’ Biometrika 58, 525-534.

Remington D.L., Leinonen P.H., Leppala J. & Savolainen O. (2013) Complex genetic effects on early vegetative development shape resource allocation differences between Arabidopsis lyrata populations. Genetics 195, 1087-1102.

Sletvold N. & Agren J. (2015) Climate-dependent costs of reproduction: Survival and fecundity costs decline with length of the growing season and summer temperature. Ecology Letters 18, 357-364.

T??rj??k O., Witucka-Wall H., Meyer R.C., Von Korff M., Kusterer B., Rautengarten C., … Altmann T. (2006) Segregation distortion in Arabidopsis C24/Col-0 and Col-0/C24 recombinant inbred line populations is due to reduced fertility caused by epistatic interaction of two loci. Theoretical and Applied Genetics 113, 1551-1561.

Tisne S., Serrand Y., Bach L., Gilbault E., Ben Ameur R., Balasse H., … Loudet O. (2013) Phenoscope: An automated large-scale phenotyping platform offering high spatial homogeneity. Plant Journal 74, 534-544.

Xu E., Vaahtera L., Hõrak H., Hincha D.K., Heyer A.G. & Brosche M. (2015) Quantitative trait loci mapping and transcriptome analysis reveal candidate genes regulating the response to ozone in Arabidopsis thaliana. Plant, Cell and Environment 38, 1418–1433.

Xu M.Y., Zhang L., Li W.W., Hu X.L., Wang M.B., Fan Y.L., … Wang L. (2014) Stress-induced early flowering is mediated by miR169 in Arabidopsis thaliana. Journal of Experimental Botany.

